# All hands on deck: Large-scale (re)sculpting of cortical circuits in post-resection children

**DOI:** 10.1101/2020.07.01.183400

**Authors:** Anne Margarette S. Maallo, Michael C. Granovetter, Erez Freud, Sabine Kastner, Mark A. Pinsk, Christina Patterson, Marlene Behrmann

**Affiliations:** Department of Psychology and Neuroscience Institute, Carnegie Mellon University; School of Medicine, University of Pittsburgh; Department of Psychology and The Centre for Vision Research, York University; Princeton Neuroscience Institute, Princeton University; Department of Psychology, Princeton University; Department of Pediatrics, University of Pittsburgh

**Keywords:** Functional connectivity, plasticity, epilepsy, lobectomy, hemispherectomy

## Abstract

Despite the relative successes in the surgical treatment of pharmacoresistant epilepsy, there is rather little research on the neural (re)organization that potentially subserves behavioral compensation. Here, we examined the post-surgical functional connectivity (FC) in children and adolescents who have undergone unilateral cortical resection and, yet, display remarkably normal behavior. Conventionally, FC has been investigated in terms of the mean correlation of the BOLD time courses extracted from different brain regions. Here, we demonstrated the value of segregating the voxel-wise relationships into mutually exclusive populations that were either positively or negatively correlated. While, relative to controls, the positive correlations were largely normal, negative correlations among networks were increased. Together, our results point to reorganization in the contralesional hemisphere, possibly suggesting competition for cortical territory due to the demand for representation of function. Conceivably, the ubiquitous negative correlations enable the differentiation of function in the reduced cortical volume following a unilateral resection.

## 1 Introduction

Accumulating evidence has shown that surgical resection can be more efficacious than pharmacological therapy in the management of drug-resistant epilepsy (Yoo & Panov, 2019; Widjaja et al., 2020). While there are promising outcomes in surgical cases involving adults (Ichikawa et al., 2020), referral for surgery evaluation in earlier stages of epilepsy is increasingly encouraged (Picot et al., 2016, Holm et al., 2018, Baud et al., 2018). One recent study (Helmstaedter et al., 2020) with a large cohort of children reported that, at one year post-surgery, anywhere from 21-50% of the individuals showed improvement in at least one of the following domains: motor function, attention, verbal memory, figural memory, language, visuoconstruction, IQ, and behavior, and with individual significant gains in the 16-42% range. This study clearly suggests that resective surgery, especially earlier in life, may permit normal development.

Consistent with these reassuring results, children with a unilateral resection of left or right ventral occipito-temporal cortex (VOTC) exhibit normal microstructural properties of the major white matter pathways in the contralesional VOTC (Maallo et al., 2020). Additionally, functional magnetic resonance imaging (fMRI) studies of such cases revealed normal activation profiles in response to different visual categories (e.g. faces and places) in the preserved cortical areas (Liu et al., 2018, Liu, Freud et al., 2019), as determined by the magnitude of selectivity, number of voxels, spatial organization of regions of interest (ROIs), and representational similarity analysis. It can be inferred from these studies that the homotopic regions in the preserved hemisphere assume the cognitive load of the resected tissue and that this occurs without any obvious cost (‘crowding’, Teuber 1975) to the overall cortical functional capacity (Beharelle et al., 2010; Wilke et al., 2009). The broader question, then, is how the (sometimes, drastically) reduced cortical territory accommodates the functions that enable typical overt behaviors.

### Underlying mechanism supporting positive post-surgical outcomes

Plasticity is an intrinsic property of the central nervous system and, although much is known about the factors that influence plasticity such as the nature, severity, and extent of the insult, as well as developmental stage, cognitive capacity, gender, genetics and access to rehabilitation (Anderson et al., 2011), identifying a candidate neural process supporting — or even driving – the changes continues to be elusive. One possible mechanism that might reflect post-surgical plasticity is changes in functional connectivity (FC), or altered correlations between the time series of different, preserved brain regions.

Many studies have documented altered post-surgical FC among networks or ROIs. For example, Morgan et al. (2019) reported that post-surgical FC, especially that of thalamus and hippocampus, may contribute to long-term seizure outcome and Liao et al. (2016) revealed differences in FC in patients, especially in the temporoparietal junction and its connection with the ventral prefrontal cortex. Typical language localization after surgery has been associated with better functional integration of various networks including the default mode network (Ibrahim et al., 2015) and Ivanova et al. (2017) described at least partial preservation of FC of language areas in the intact right hemisphere. Last, relative to controls, Kliemann et al. (2019) uncovered increased FC between different networks (e.g., default and somatomotor networks), but not within-network, in the preserved hemisphere of adults with hemispherectomy (sometimes referred to as ‘hemispherotomy; see Kim et al., 2018), and, provocatively, suggested that this increased connectivity might compensate for cognitive impairments. It still remains to be determined, however, exactly how these connectivity changes contribute to the surprising recovery of cognitive abilities in many post-surgical patients (Uddin, 2020).

### Characterizing detailed functional connectivity profiles

In light of the FC changes post-surgically, and the greater potential for plasticity during childhood than adulthood (Anderson et al., 2011; Helmstaeder et al., 2020; Lidzba et al., 2017), we undertook to characterize comprehensively the FC in the preserved hemisphere of a pediatric sample. We used data acquired during a visual category localizer task, rather than during resting state, both to ensure that we were able to elicit sufficient BOLD signal from the young patients and to validate the connectivity of functionally specific ROIs. We included a closely age-matched control group, as FC becomes more integrated across development, with the brain shifting from more localized to more global patterns (Vohryzek et al., 2020, Puxeddu et al., 2020). We analyzed only the preserved hemisphere in patients, given that heterotopic, intrahemispheric connectivity changes are more likely to be plastic compared with stable homotopic connections which have direct anatomical projections (Shen et al., 2015).

Inspired by Kliemann et al. (2019), we too compared FC between- and within-networks in patients and controls. However, rather than focus on functionally coupled networks, we examined correlations at two parcellation levels: among anatomically demarcated ROIs, without assumptions about regional coupling, and among networks comprised of subgroups of anatomical ROIs.

Last, we examined the effect of structural distance on FC in the patients versus controls. An efficient cortex depends on both long- and short-range connections: whereas long-range FC requires increased time and metabolic costs (Bullmore & Sporns, 2012; Liang et al., 2013), short-range FC typically commands less time and metabolic costs, and exhibits stronger FC strength (Salvador et al., 2005). In an assumption-free fashion, we compared FC across short, intermediate and long distances (see Achard et al., 2006; He et al., 2007). In addition to the bottom-up parcellation and the distance-scaled approaches, in contrast with most studies that focus on average correlations independent of sign, in all analyses, we separated the FC into positive and negative correlations.

## 2 Results

We present findings from a group of nine children with unilateral cortical resection (Table 1) and nine controls. The patients exhibited largely normal visual perceptual abilities; even if one measure were atypical, other measures were normal (Supp. Mat. Table S1; Liu, Freud et al., 2019).

**Table 1.** Patient history. Ages at scan and at surgery are in years, except for SN.

| Patient Code* | Sex | Age, scan | Age, surgery | Surgical procedure | Engel Epilepsy<br>Surgical Outcome* |
| --- | --- | --- | --- | --- | --- |
| *Right focal resection* |  |  |  |  |  |
| EK | M | 17 | 17 | Gross total resection of right frontal neuroglial/gliotic lesion | IA |
| KQ | F | 16 | 15 | Right anterior temporal lobectomy and hippocampectomy | IA |
| UD | M | 13 | 6 | Right occipital and posterior temporal lobectomy with resection of inferomesial temporal dysembryoplastic neuroepithelial tumor | IA |
| *Left focal resection* |  |  |  |  |  |
| DX | M | 16 | 15 | Left frontal corticectomy with resection of focal cortical dysplastic lesion | IA |
| NN | M | 18 | 15 | Left occipital lobectomy, left posterior temporal and left parietal corticectomy with extant polymicrogyria | IIA |
| SN | M | 15 | 1day | Evacuation of left temporal hematoma | IA |
| TC | F | 15 | 13 | Left parietal and occipital lobectomy | IID |
| *Left hemispherectomy* |  |  |  |  |  |
| FD | M | 14 | 7 | Left frontal and parietal lobectomy followed by left anatomic hemispherectomy | IID |
| JF | M | 16 | 5 | In utero bilateral stroke, left hemispherectomy | IA |

\*The initials used here are codes used to protect the identity of participants and are not the patients' true initials.
\*For details of the Engel outcome scale, see Engel (1993).

We hypothesized that the FC of the contralesional hemisphere might be altered to accommodate the functions of the resected tissue. For each analysis, we first report the profile of the group of control subjects and then of the group of patients. We then compare the two groups and follow-up by evaluating each patient individually against the control distribution. At the end of this section, we provide a summary of all the analyses with the main results, along with the relevant figures and/tables.

### 2.1 FC between- and within-regions of interest

Our first analyses were done at the level of functionally defined ROIs in which we used a univariate approach to localize category-specific ROIs, which were preferentially responsive to different categories (e.g. faces). In addition, given that FC changes have been reported for subcortical structures (Morgan et al., 2019), we also included two anatomically demarcated vision-relevant subcortical ROIs: the lateral geniculate nucleus (LGN) and the pulvinar (see Section 4.4 for details). In the next analysis of anatomically defined ROIs, we used affine and non-linear transformations to register the HCP atlas and get anatomical parcellations in each participant’s native volumetric space (see Section 4.5).

Procedurally, we extracted the evoked BOLD signal time series from every grey matter voxel (following anatomical segmentation) comprising each ROI in the controls’ left and right hemispheres (LH and RH, respectively) and in patients’ contralesional hemispheres (see Sections 4.6 for details). Here, for each ROI pair, we used the mean of all the voxel-to-voxel correlations as the main measure for FC.

#### 2.1.1 Voxel-wise FC across functional ROIs

Because our paradigm evoked BOLD signals in response to images, we analyzed FC at the category-selective ROI level (selective for different categories: faces, objects, places, words, scrambled objects) and for the LGN and pulvinar. In controls, we were unable to identify a word-selective region or the LGN in the RH, for 6/9 and for 1/9, respectively. We averaged the FC (Supp. Mat. Fig. S2A, E) and examined relationship of the between- and within-ROI FC. Conventionally, FC has been characterized as the mean of the correlations of all the BOLD time courses extracted from the ROIs (at the voxel or vertex level for volumetric or surface analysis). Because both positive and negative correlations have been reported and the supposition is that they play different roles, we separated the FC results by the sign of the correlation.

Upon inspection, we confirmed that there were mutually exclusive pairs of voxels that were either positively or negatively correlated among all ROI pairs. Notably, there were rather few negative correlations among the category cortical ROIs, especially within the same functional ROIs (along the diagonal, Supp. Mat. Fig. S2B, F). This profile is intuitive given that category-specific signals, which ought to be highly correlated during a visual task, are positively co-evoked in the ROIs. Interestingly, the fraction of negative correlations in the subcortical category ROIs differed from the cortical category ROIs: for the LGN, the fraction of negative correlations (light blue to cyan, Supp. Mat. Fig. S2A, E) was not as low as in the cortical category ROIs; for the pulvinar, the fraction of negative correlations (green, Supp. Mat. Fig. S2A, E) was roughly 50%. Also, the LGN showed higher FC with the cortical category ROIs compared to the pulvinar, suggesting that the former is more functionally specific than the latter.

We proceeded to study the effects of the different factors affecting the combined, positive, and negative (between-ROI only; no negative within-ROI) FC, separately in patients and controls. First, in controls, we performed a two-way ANOVA on FC, with hemisphere (LH versus RH) and ROI connectivity type (between versus within-category ROI) as the factors, and found no significant interaction effects (combined FC: F(1, 32) = 1.28, p = 0.2666; positive FC: F(1, 32) = 0.41, p = 0.5274), and no significant main effects of hemisphere (combined FC: F(1, 32) = 0.15, p = 0.7013; positive FC: F(1, 32) = 0.78, p = 0.3839). There was a significant main effect of ROI connectivity type on the combined FC (F(1, 32) = 793.89, p < 0.001) where the between-category ROI FC magnitude (mean: 0.0517) was smaller than the within-ROI FC (mean: 0.2782), as well as on the positive FC in the same direction ((F(1, 32) = 706, p < 0.001); between-ROI mean (0.1154) < within-ROI mean (0.3010)). Given that there were no negatively correlated voxels within the same category ROI, we performed a paired t-test on the LH’s and RH’s between-category ROI negative FC, and found no significant interhemispheric differences (|t| = 2.1071, p = 0.062, df = 8).

Next, in patients, we localized all category-selective ROIs in the contralesional hemisphere, but some ROIs were missing in some patients: EK: no LH object-selective, KQ: no LH word-selective, UD: no LH word-selective, FD: no RH place or word-selective, also no subcortical ROIs, JF: no RH object-selective. Nevertheless, intermediate and high-level visual behaviors were largely comparable to those of controls (Supp. Mat. Table S1).

As with controls, we computed the voxel-wise correlation in functional ROIs from the patients’ contralesional hemisphere, averaged over the three patients with right resection and the six patients with left resection (Supp. Mat. Fig. S2I, M, respectively). A paired t-test revealed that the between-category ROI FC was always smaller in magnitude than the within-category ROI FC (combined FC: |t| > 18.4, p <0.001, df = 8; positive FC: |t| > 17.4, p < 0.001, df = 8). As in controls, the number of negative voxel-wise correlations in patients within the same (cortical) functional ROIs was essentially zero, while the subcortical functional ROIs exhibited a graded positive/negative split (Supp. Mat. Fig. S2J, N).

An unbalanced two-way ANOVA on the combined and positive FC, separately, with group (patients versus controls) and ROI connectivity type (between versus within) as factors revealed no significant interactions between group and ROI connectivity type (combined FC: F(1, 50) = 0, p = 0.9683; positive FC: F(1, 50) = 0.04, p = 0.8336), and no significant main effects of group (combined FC: F(1, 50) = 1.92, p = 0.1723; positive FC: F(1, 50) = 0.95, p = 0.3342). Because there was no within-ROI negative FC in patients, we used a two-sample t-test on the between-ROI FC on the two groups and found no significant group differences (|t| = 1.0587, p = 0.2998, df = 25). However, there was still a significant main effect of ROI connectivity type on (1) combined FC (F(1, 50) = 880.23, p < 0.001), between-ROI mean (0.0481) < within-ROI mean(0.2747)); and (2) positive FC (F(1, 50) = 879.26, p < 0.001), between-ROI mean (0.1129) < within-ROI mean (0.2994)).

Finally, using the modified t-test (Crawford & Howell, 1998) for single-subject comparisons of patient data and the controls, with correction for multiple comparisons, only two patients, NN and JF, fell outside the control distribution for either ROI connectivity type or FC Index (Supp. Mat. Table S3).

In sum, the results from FC of visual ROIs revealed that, for both control and patient groups, the between-category ROI FC was weaker than the within-category ROI FC, be it in the combined or positive FC, and that the FC of LH in controls was the same as the FC of the RH (except in NN, with extant polymicrogyria and low IQ pre-surgically; and in JF, with a very large resection). These results are consistent with previous findings of normal neural responses and category-selectivity (Liu et al., 2018, Liu, Freud et al., 2019). The normal correlation profiles of the patients serve as validation that the data quality is sound and that we can replicate the expected result in which cortical regions that serve the same function are positively co-activated.

#### 2.1.2 Voxel-wise FC across anatomical ROIs

With the above validation in hand, we computed the FC among all pairs of anatomically-defined ROIs in controls (Fig. 1A, E, LH and RH. respectively). We first determined whether there were any hemispheric asymmetries and then evaluated differences within-ROI FC (mean of diagonals, Fig. 1A, E) and between-ROI FC (mean of off-diagonals, Fig. 1A, E). We then performed a two-way ANOVA on FC, with hemisphere (LH/RH) and ROI connectivity type (between-/within-ROI) as the factors. The results showed no main effect of hemisphere (F(1, 32) = 0.47, p = 0.4988) and no significant interaction (F(1, 32) = 0.16, p = 0.6946). There was, however, a main effect of ROI connectivity type (F(1, 32) = 211.77, p < 0.001), with between- (mean: 0.0226) smaller than within-ROI FC (mean: 0.1287).

**Figure 1.**
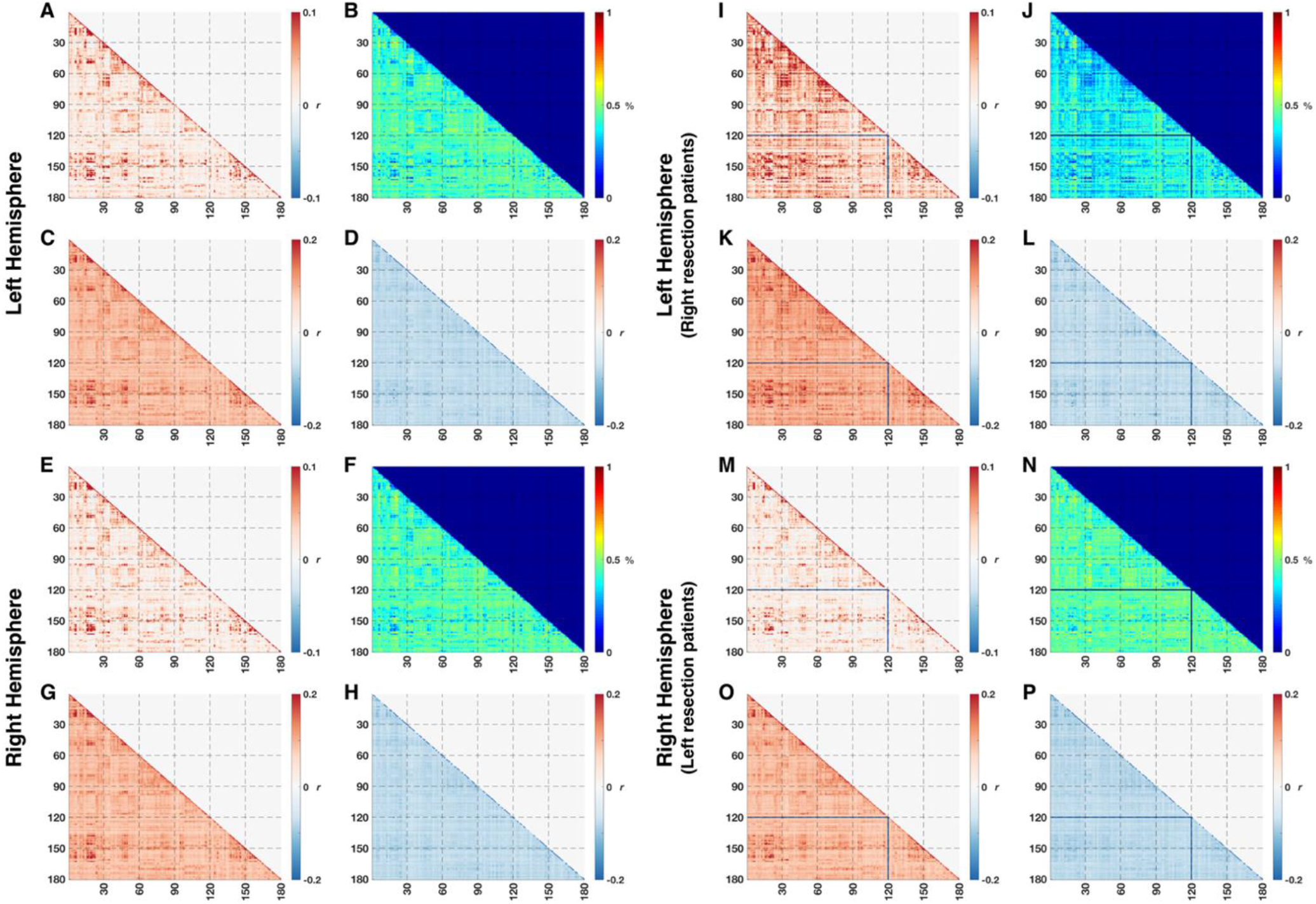
FC among 180 ROIs spanning the entire hemisphere averaged over (A-H) controls or over (I-L) three patients with right resection or (M-O) six patients with left resection. Diagonal values (within-ROI FC) are larger in absolute value than off-diagonal values (between-ROI FC, c.f. Supp. Mat. Table S2). (A, E, I, M) FC were averaged over all possible voxel-to-voxel pairs from corresponding pairs of ROIs. (B, F, J, N) Percentage of voxel-to-voxel connection in each ROI pair that are negatively correlated is roughly 50% while the remaining fraction is positively correlated. FC average over voxelwise connections with only (C, G, K, O) positive or (D, H, L, P) negative correlations. Note the different color scales between A/E/I/M and C/D/G/H or K/L/O/P, indicating stronger FC values after splitting into purely positive or purely negative values. For ROI labels, see (Glasser et al., 2016). Lines in patients’ matrices indicate one ROI (hippocampus) that was not identifiable in patients EK and TC.

When we examined the positive and negative correlations separately, we discovered that the positive/negative split was at roughly 50%/50% (Fig. 1B and F, respectively). After separating the FC into positive (Fig. 1C, G) or negative (Fig. 1D, H) correlations, correlations had greater absolute values than when averaged. To determine the relationship of between- and within-ROI FC for the independent positive or negative FC from control data, we performed a two-way ANOVA (hemisphere, ROI connectivity type) separately, and found no significant interactions (positive FC: F(1, 32) = 0.11, p = 0.7383; negative FC: F(1, 32) = 0.04, p = 0.8427) and no main effects of hemisphere (positive FC: F(1, 32) = 0.38, p = 0.5416; negative FC: F(1, 32) = 0.18, p = 0.6733). However, there was a significant main effect of ROI connectivity type on FC, with the magnitude of between-ROI mean being smaller than the magnitude of within-ROI mean for both positive (F(1, 32) = 227.66, p < 0.001, between-ROI mean: (0.0962) < within-ROI mean(0.1890)) and negative (F(1, 32) = 4.91, p = 0.034, between-ROI mean(|-0.0780|) < within-ROI mean (|-0.0814|)) FC. These results indicate that the magnitude of between-ROI FC was consistently lower than within-ROI FC, be it in the combined, positive, or negative correlations and that the FC of LH and RH in controls were comparable.

We then computed the voxel-wise correlation in the patients’ contralesional hemisphere, averaged over the three patients with RH resection and the six patients with LH resection (Fig. 1I, M, respectively). A paired t-test (between-versus within-) revealed that between- (mean: 0.0259) was smaller than within-ROI FC (mean: 0.1312) (|t| = 17.1463, p < 0.001, df = 8), similar to the result from controls. As in controls, some correlations were negative (Fig. 1J, N), and we separated the correlations into positive (Fig. 1K, O) or negative (Fig. 1L, P) correlations. Paired t-tests confirmed that the magnitude of between-was smaller than within-ROI FC for both positive (|t| = 20.4584, p < 0.001, df = 8, between-ROI mean (0.0997) < within-ROI mean (0.1934)) and negative FC (|t| = 3.4535, p = 0.0086, df = 8), between-ROI mean(|-0.0790|) < within-ROI mean (|-0.0848|)).

Next, we performed an unbalanced two-way ANOVA, separately, on the combined, positive, and negative FC, with group (patients versus controls) and ROI connectivity type (between versus within-ROI) as factors. Given that there were no significant differences between controls’ LH and RH above, we compared the contralesional hemisphere from patients to the combined (concatenated) data from LH and RH in controls. There was no interaction between group and ROI connectivity type (combined FC: F(1, 50) = 0, p = 0.951; positive FC: F(1, 50) = 0.48, p = 0.8281; negative FC: F(1, 50) = 0.71, p = 0.4044), and no main effect of group (combined FC: F(1, 50) = 0.23, p = 0.63; positive FC: F(1, 50) = 0.48, p = 0.4903; negative FC: F(1, 50) = 2.52, p = 0.1188), indicating equal FC for patients and controls. However, the main effect of ROI connectivity type was significant on (1) combined FC (F(1, 50) = 307.64, p < 0.001), between- (0.0237) < within-ROI mean (0.1295); (2) positive FC (F(1, 50) = 387.02, p < 0.001), between- (0.0970) < within-ROI mean (0.1905); and (3) negative FC (F(1, 50) = 10.21, p = 0.0024), between- (|-0.0783|) < within-ROI mean (|-0.0825|).

Last, the modified t-test (Crawford & Howell, 1998) for comparing individual patients to the distribution of control data from the corresponding hemisphere revealed no significant differences in any individual patient com for either ROI connectivity type (or for FC Index, which is the ratio of the between-to within-ROI FC, Kliemann et al., (2019)), correcting for multiple comparisons (Supp. Mat. Table S4).

These results repeatedly showed no significant differences in FC between patients and controls, at the level of anatomical ROIs. The important conclusion is that the magnitude of between-ROI FC was always smaller than the magnitude of the within-ROI FC in both groups, revealing stronger connectivity among voxels belonging to the same than to different ROIs. This finding is intuitively consistent with the high-level notion that voxels (thereby, presumably, neurons) from a specific cortical region are co-activated and engaged in particular ensembles together. This is also consistent with the finding from the functionally defined category ROIs (Section 2.1.1) in which the magnitude of within-ROI positive FC was larger in the (cortical) functional ROIs. However, in contrast to the FC of the cortical functional ROIs, there was an even split into positive or negative correlations within the same anatomical ROIs. Concurrently, the positive FC in the category ROIs was larger than the positive FC in the anatomical ROIs. Together, these results uncover the stronger correlations in task-related regions (visual cortex/visual stimulation) and the roughly equal proportion of positively and negatively correlated voxels in the anatomical ROIs may be a hallmark of to the non-specificity of anatomical ROIs in response to the visual task.

### 2.2 FC between different and within the same networks

Next, we compared the FC of the 22 HCP atlas networks (Supp. Mat. Table S5, see also Glaser et al., 2016) between patients and controls, with each network consisting of a subgroup of the 180 anatomical ROIs presented above. The procedures here mirrored those used in the previous section, only with the ROIs replaced by networks (see Sections 4.5 and 4.6 for details).

#### 2.2.1 Voxel-wise FC across networks

Graphically, in the controls, the results were largely comparable in magnitude to that of Kliemann et al. (2019), with values in the range of |r|~0.1 in both hemispheres (Fig. 2A, E), with minor differences. Kliemann et al. (2019) z-transformed their correlation coefficients, but we present the data in raw correlation values, given that at the range of values here, a z-transformation would be strictly linear and would not drastically alter the values.

**Figure 2.**
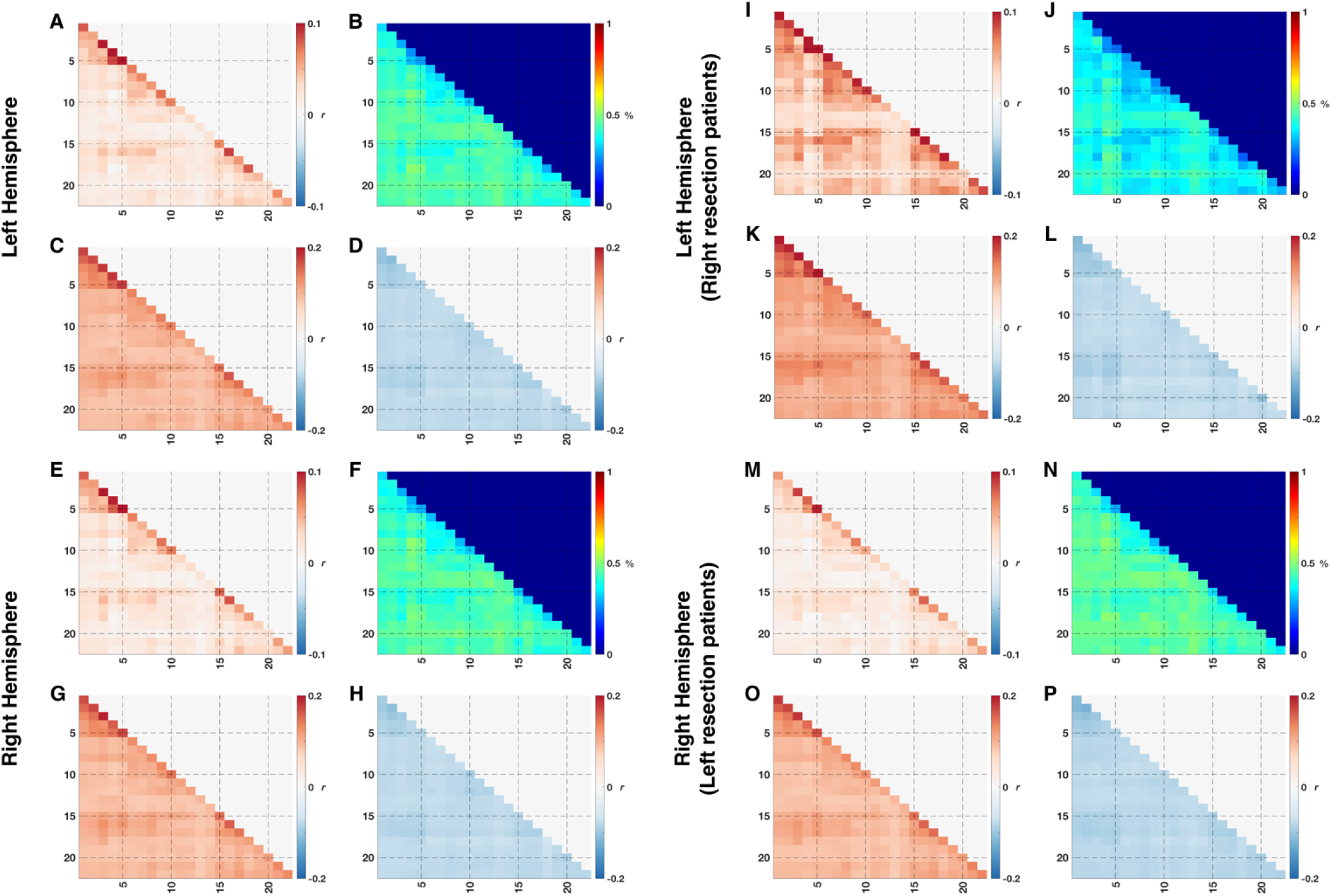
FC among 22 networks averaged over (A-H) controls or over (I-L) three patients with right resection or (M-O) six patients with left resection. Diagonal values (within-network FC) are larger in absolute value than off-diagonal values (between-network FC, c.f. Supp. Mat. Table S5). (A, E, I, M) FC were averaged over all possible voxel-to-voxel pairs from corresponding pairs of networks. (B, F, J, N) Percentage of voxel-to-voxel connection in each network pair that are negatively correlated is roughly 50% while the remaining fraction is positively correlated. FC was averaged over voxel-wise connections with only (C, G, K, O) positive or (D, H, L, P) negative correlations. Note the different color scales between A/E/I/M and C/D/G/H or K/L/O/P, indicating stronger FC values after splitting into purely positive or purely negative values. For network labels, see Supp Materials Table S7.

To illustrate the relationship of within-network (mean of diagonals, Fig. 2A, E) and between-network FC (mean of off-diagonals, Fig. 2A, E) in controls, we performed a two-way ANOVA on FC, with hemisphere (LH versus RH) and network connectivity type (between-versus within-network) as factors. There were no significant interactions (F(1, 32) = 0.07, p = 0.7864) and no significant main effect of hemisphere (F(1, 32) = 0.29, p = 0.5932), but there was a significant main effect of network connectivity type (F(1, 32) = 107.89, p < 0.001), with the between-network FC (mean: 0.0212) being smaller than the within-network FC (mean: 0.0623).

Similar to the splitting into positively or negatively correlated voxels in the HCP anatomical ROIs, we found a roughly 50%/50% splitting of the voxels comprising the network pairs (Fig. 2B, F). From a two-way ANOVA on positive and negative FC (hemisphere, network connectivity type), there were no interactions (positive FC: F(1, 32) = 0.07, p = 0.7861; negative FC: F(1, 32) = 0.02, p = 0.8811), no significant main effects of hemisphere (positive FC: F(1, 32) = 0.52, p = 0.476; negative FC: F(1, 32) = 0.75, p = 0.3917), but there was still a significant main effect of network connectivity type (positive FC: F(1, 32) = 145.78, p < 0.001; negative FC: F(1, 32) = 21.46, p = 0.0001), where the magnitude of the between-network FC (positive mean: 0.0957, negative mean: −0.0784) was smaller than the magnitude of the within-network FC (positive mean: 0.1374, negative mean: −0.0834).

The results from the network-level analysis support the claim that the roughly equal balance of positive and negative correlations is due to the non-specificity (of the networks) in the visual task. Nevertheless, to ensure that the negative correlations were not simply an artefact of the processing pipeline as negative correlations might arise from regression of nuisance signals (Murphy et al., 2009), these analyses were redone in several ways: by separately regressing the mean signal from white matter only, or the cerebrospinal fluid only, or by regressing global signal alone and, then, by regressing various combinations of the above. The results appear unchanged by these various manipulations – that is, the positive/negative split was evident in all preprocessing combinations, even in minimally processed data, which were only volume-registered and co-registered to the anatomical image without any nuisance signal regression (findings of these analyses in Supplementary Figure S6).

Next, for networks in the patients’ contralesional hemispheres, we computed the voxel-wise correlation averaged over the three patients with RH resection and the six patients with LH resection (Fig. 2I, M, respectively). A paired t-test confirmed that the between-network FC (mean: 0.0246) was smaller than the within-network FC (mean: 0.0650) (|t| = 14.9468, p < 0.001, df = 8), similar to the result from controls. Also, as in controls, there was a similar division into positive and negative correlations (Fig. 2J, N). After separating the correlations into positive (Fig. 2K, O) or negative (Fig. 2L, P) correlations, paired t-tests revealed that the magnitude of between-was smaller than within-network FC for both positive (|t| = 20.1991, p < 0.001, df = 8), between- (0.0996) < with-network mean (0.1417)) and negative FC (|t| = 7.604, p < 0.001, df = 8), between- (|-0.0796|) < with-network mean (|-0.0855|)).

Again, we performed an unbalanced two-way ANOVA on the combined, positive, and negative FC, separately, with group (patients versus controls) and network connectivity type (between versus within-network) as factors. There were no significant interactions between group and network connectivity type (combined FC: F(1, 50) = 0.01, p = 0.9139; positive FC: F(1, 50) = 0, p = 0.9473; negative FC: F(1, 50) = 0.18, p = 0.6696) and no significant effect of group (combined FC: F(1, 50) = 0.62, p = 0.4342; positive FC: F(1, 50) = 1.82, p = 0.1834; negative FC: F(1, 50) = 2.35, p = 0.132). However, there was still a significant main effect of network connectivity type on (1) mean FC (F(1, 50) = 111.59, p < 0.001), between- (0.0223) < within-network mean (0.0632)); (2) positive FC (F(1, 50) = 193.38, p < 0.001), between- (0.0970) < with-network mean (0.1388)); and (3) negative FC (F(1, 50) = 24.55, p < 0.001), between- (|-0.0788|) < within-network mean (|-0.0841|)).

Last, single-subject comparisons (Crawford & Howell, 1998) uncovered no significant differences in any patient compared to the control distribution for either network connectivity type after correcting for multiple comparisons (Supp. Mat. Table S7).

#### 2.2.2 Altered negative FC in patients at the network-level

Thus far, the FC averaged over all the different ROI/network pairs revealed no group or single-subject differences in the between- or within-ROI/network FC. It is possible, however, that there may be *isolated network pairs* that differ between the groups but, because we only examined the means (over a large number of between- or within-network FC), this was not apparent. We thus compared each single network pair FC of individual patients to the corresponding network pair FC in controls (see Section 4.7 for details). We considered a particular network pair to have high variability in FC if it exceeded a distance of two standard deviations from the control mean (FC > 2 standard deviations from the mean, criterion 1). Furthermore, because such variability might spuriously arise from intrinsic individual differences and rather than from a patient’s resection, we conducted a leave-one-out test on the control data to determine the (maximum) number of the network pairs (criterion 2) that exhibited high variability. Interestingly, there were fewer network pairs that exhibited high variability in controls’ negative FC for either the LH (up to 75) or RH (up to 49) than in their positive FC (LH: up to 158, RH: up to 168), indicating a tendency of the negative FC to be relatively stable in controls, a point we return to in the Discussion.

In patients, several network pairs had positive FC values (Fig. 3A-C) that deviated from the control mean by more than two standard deviations (satisfied criterion 1), especially in patients with focal RH resection (see Figure 3; abundance of red cells in EK and UD, specifically, indicating larger positive FC than controls). However, the number of pairs that deviated from the control mean was under the maximal number of high variability network pairs in the corresponding hemisphere in controls (values from criterion 2 in patients were smaller than controls’). For instance, in positive FC in the LH, patients UD and KQ deviated in 138 and 114 network pairs, respectively, fewer than the 158 network pairs showing high variability in controls. Therefore, these high FC might simply be due to intrinsic individual differences, and thus we concluded that all patients exhibited normal positive FC.

**Figure 3.**
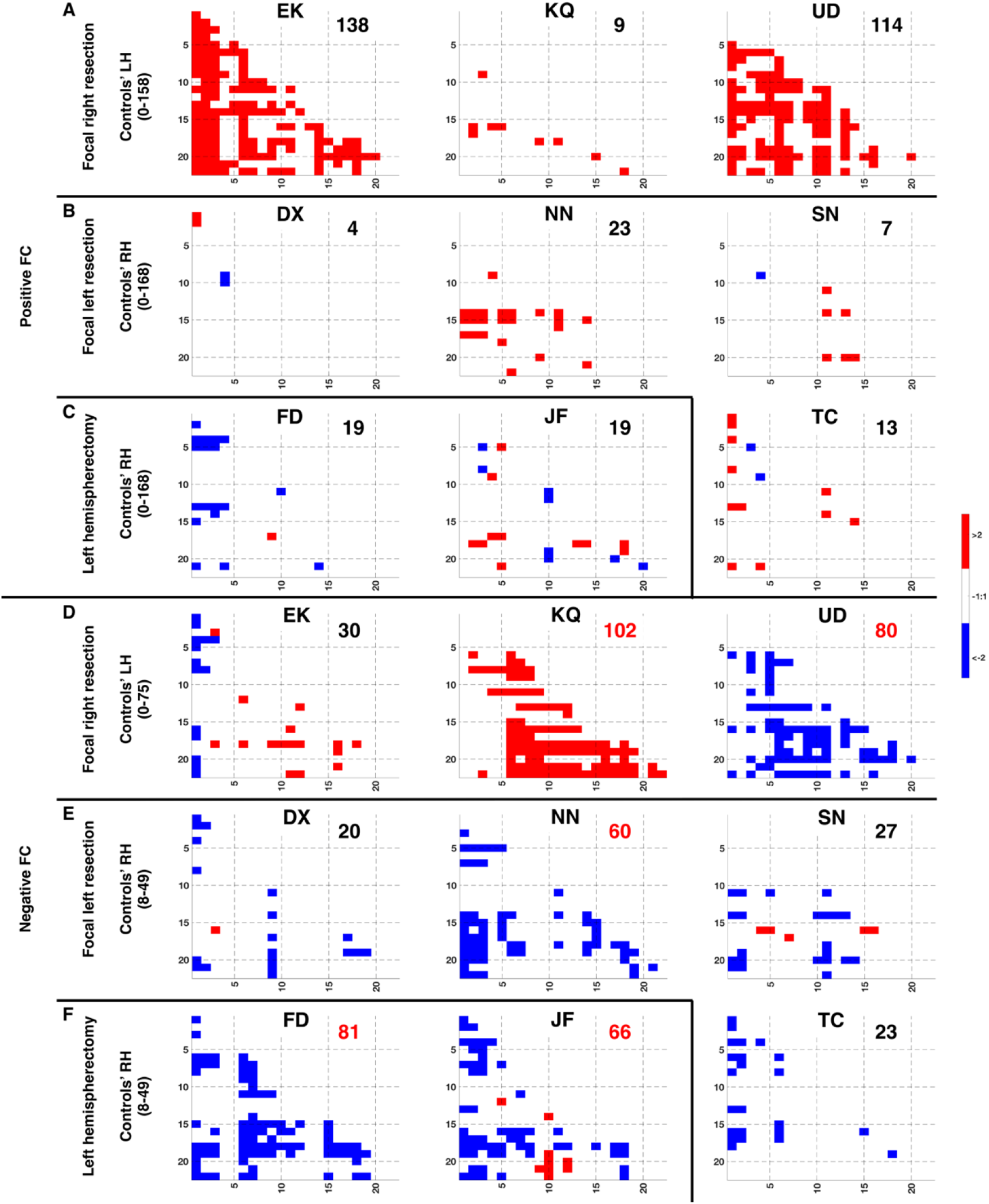
Distance-from-mean in (A-C) positive or (D-F) negative FC of patients compared to controls. Red or blue cells indicate FC with network pairs that are more than two standard deviations greater or lower, respectively, than the control mean (satisfied criterion 1). There are more network pairs (simply the count of red/blue cells, criterion 2) that are different between patients (KQ, UD, NN, FD, and JF) and controls only in the negative FC. Values for controls are number of network pairs that exhibited variability in individual controls. Inset numbers in each patient’s matrices are number of network pairs that are more than two standard deviations away from controls’ mean in either direction. Numbers in red exceed the maximum variable network pair numbers in controls.

In contrast, five patients (KQ, UD, NN, FD, JF) had a significantly larger number of high variability network pairs in the negative FC (Fig. 3D-F). For example, KQ’s LH had 102 network pairs, greater than the max 75 pairs in controls LH, as identified in the leave-one-out test, and FD had 80 networks-pairs that deviated from the max 49 pairs in control RH. Curiously, KQ had negative FC values that were smaller in magnitude than controls (Fig. 3D: red cells indicate FC that was more than two standard deviations higher than controls; i.e. less negative than controls), while UD, NN, FD, and JF had negative FC that were larger in magnitude than controls (Fig. 3D-F: blue cells indicate FC that was more than two standard deviations lower than controls; i.e. more negative than controls). The remaining patients (EK, DX, SN, TC) had some network pairs that deviated from the control mean, but the number of such network pairs in each patient was smaller than the maximal value in controls.

In sum, the mere differences in positive FC in patients and controls could be attributed to intrinsic individual differences, owing to the comparable variability seen in healthy, typically developing controls. Nevertheless, patients exhibited high variability in more network pairs than controls in the negative FC (and this was independent of which hemisphere was preserved).

#### 2.2.3 Networks that are stable in controls are altered in patients with resection

In this next analysis, we determined whether there were *specific network pairs* that differed between the groups. To this end, we determined the overlap of the distribution of the network pairs with high variability in controls, and only focused in patients on network pairs for which no control exhibited variability. Fig. 4 shows a heatmap of controls showing which network pairs were highly variable (gray cells, Fig. 4) and which were highly stable (black cells under diagonal, Fig. 4). This was derived by counting how many controls, out of nine, deviated from the remaining controls’ mean by more than two standard deviations, in the leave-one-out analysis (see Section 4.8 for details). The gray cells in the control heatmap – indicating that at least one control deviated from the rest of the group – occupied as much as 84% of the entire LH or 89% of the RH in the positive FC heatmaps, and as much as 81% of either hemisphere in the negative FC heatmaps, although, the absolute locations of the gray cells were variable for each configuration (e.g. compare gray heat map of LH positive versus RH negative, Fig. 4). That these differences were random and mostly non-overlapping confirms that these are due to intrinsic individual differences. At most, there were two control participants (22% of control group) who overlapped at any given network pair in either the positive (Fig. 4A-B, gray) and negative (Fig. 4C-D, gray) FC heatmaps.

**Figure 4.**
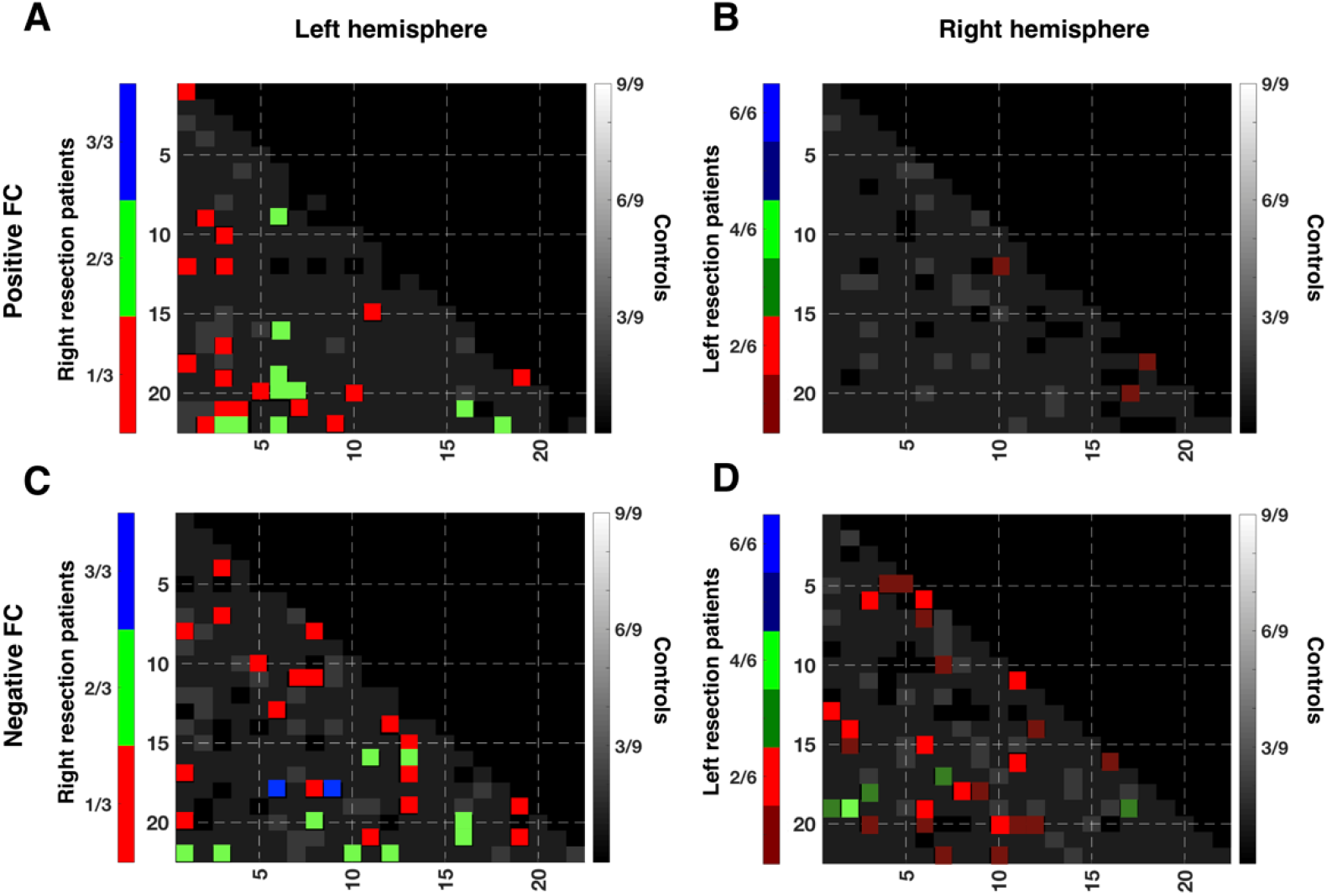
Heatmap of network pairs that show individual differences in controls and alterations to the stable network pairs in patients. Network pairs that indicate individual differences between controls are widespread and variable throughout the entire hemisphere in both (A-B, gray) positive and (C-D, gray) negative FC of either hemisphere. At most, there are 2/9 controls for whom these differences overlap at any given network pair. There are also network pairs that were stable among controls (black cells under diagonal). Differences between patients and controls in (A-B, colored) positive and in (C-D, colored) negative FC are only considered for stable network pairs where there is no variability in controls. Patient data are shown for patients with right or left resection (networks are from contralesional left or right hemisphere, respectively). Right resection patients showed consistent deviation from control mean in only two network pairs (blue cells, panel C: Posterior Cingulate cortex to Somatosensory/ Motor cortex and Posterior Cingulate cortex to Posterior opercular cortex).

At the same time, we concluded that the remaining 16%, 11% in LH/RH, respectively, in positive FC heatmap; and the remaining 19% in either hemisphere in negative FC heatmap in controls were stable, given that these network pairs did not exhibit variability in any of the controls. To determine specific network pairs in patients that were potentially susceptible to changes following a resection, we only examined these stable network pairs in patients.

We examined the networks deviation in the patients depending on the hemisphere that had been resected. Unsurprisingly, there were deviations in some of the supposedly stable network pairs and these differences were presumably a result of the pathology or the surgery (Fig.4, colored cells). No network pairs were consistently altered in all nine patients, either in positive or negative FC heatmap, and only two network pairs were commonly altered in all the three patients with RH resection (Fig. 3C, blue cells: connection between posterior cingulate cortex and: (1) somatosensory/motor Cortex; (2) posterior opercular cortex), only in the negative FC heatmap. Note however, that KQ exhibited less negative FC (smaller absolute value) while EK and UD exhibited more negative FC (i.e. larger absolute value) compared to controls. As for the six patients with LH resection, only one network pair was commonly altered in at most four patients (Fig. 4D, bright green cell: connection between early visual cortex and anterior cingulate + medial prefrontal cortex), and only in the negative FC heatmap. Nevertheless, the conclusion is that, even in the presumably stable network pairs, patients exhibited altered FC. Moreover, the probability of seeing alterations in patients among the stable network pairs was higher than the overlap of high variability network pairs in controls (e.g. green cells in Fig. 4A,C indicate two of three RH resection patients: 67% probability, or bright red in Fig. 4B,D indicate two of six LH resection patients: 33%, c.f. overlap of 22% in controls for other non-stable network pairs).

Together, these findings demonstrate that the mutually exclusive positive or negative correlations arise from the non-specificity of the response of the cortical networks to the visual stimulation, and were not simply an artefact of signal pre-processing. Importantly, by splitting the data into positive or negative FC and comparing individual patients versus controls at each single network pair, we found that all patients exhibited normal positive FC and over half of the patients had altered negative FC, even after a stringent combination of exclusion criteria. We also showed that, in controls, there were network pairs prone to variability but also network pairs that seem impervious to deviations. In contrast, patients still exhibited abnormal FC even in these latter, stable network pairs. Above all, there was no single network pair that was altered (in the contralesional hemisphere) in all RH/LH resection patients, but there were two network pairs that were altered in all the RH resection patients, in addition to other network pairs in patients that also deviated from controls.

### 2.3 Contralesional hemispheres exhibit reorganization at different distance scales

Thus far, we have observed group differences in FC both in the sign and magnitude of correlations, and both at the ROI and at the network-level. Here, without relying on *a priori* assumptions inherently conferred on the ROIs or the networks as derived from the HCP atlas, we used an assumption-free distance-scaled approach to quantify differences between patients and controls (see Section 4.9 for details), given that there can be differential alterations in FC as a function of distance, and these alterations are correlated with behavior (Diao et al., 2020). The distance-scaled measures presented here capture the level of cooperation among voxels (and, by extension, it can be argued, cortical volume) perhaps necessary to support behavior.

Briefly, for every voxel in the brain (LH and RH analyzed separately), we defined mutually exclusive communities that were delimited by linear distance: short (S), intermediate (I), and long (L) range. Any given voxel has a unique fixed population of voxels in each respective community, *S*, *I*, or *L:* this is simply the number of voxels within the pre-specified distances from the voxel of interest (see Section 4.9 for further details on distance calculation for each participant). Next, we computed a voxel’s significant correlation fraction (SCF, always a positive value) by counting the number of significant positive or negative correlations (|r |≥ 0.241, p < 0.001, df = 182), separately, that the voxel had among all the voxels in its three communities, and then dividing this count by the population sizes of the respective communities (e.g. a voxel that has 20 other voxels with which it has a positive r ≥ +0.241 in its short-range community, which comprised a total of 100 voxels, has a positive-SCF value of 0.2; or a voxel that has 5 other voxels with which it has negative correlation r ≤ −0.241 in its intermediate-range community, which comprised a total of 50 voxels, has a negative-SCF value of 0.1).

We used the 180 ROIs from each hemisphere in the HCP atlas here, only as a means to visualize and quantify the SCF on a comparable scale between patients and controls (i.e. no two participants had exactly the same number of voxels, and thus, we needed a common reference). In controls, we saw that the positive-SCF in both the LH and RH had a maximum of about 0.07 for the short community and even lower numbers for the intermediate and long communities (Fig. 5A, left panel). At the same time, the negative-SCF across all three communities were low (Fig. 5B, left panel). Specifically, in controls: (1) only a sparse volume of voxels exhibits high positively correlated co-activation; (2) the fraction of highly positively correlated voxels are diminished as the distance is increased (darker blues in long-range compared to light green in short-range, Fig. 5A); and (3) the fraction of highly negatively correlated voxels is relatively constant and low across distance.

**Figure 5.**
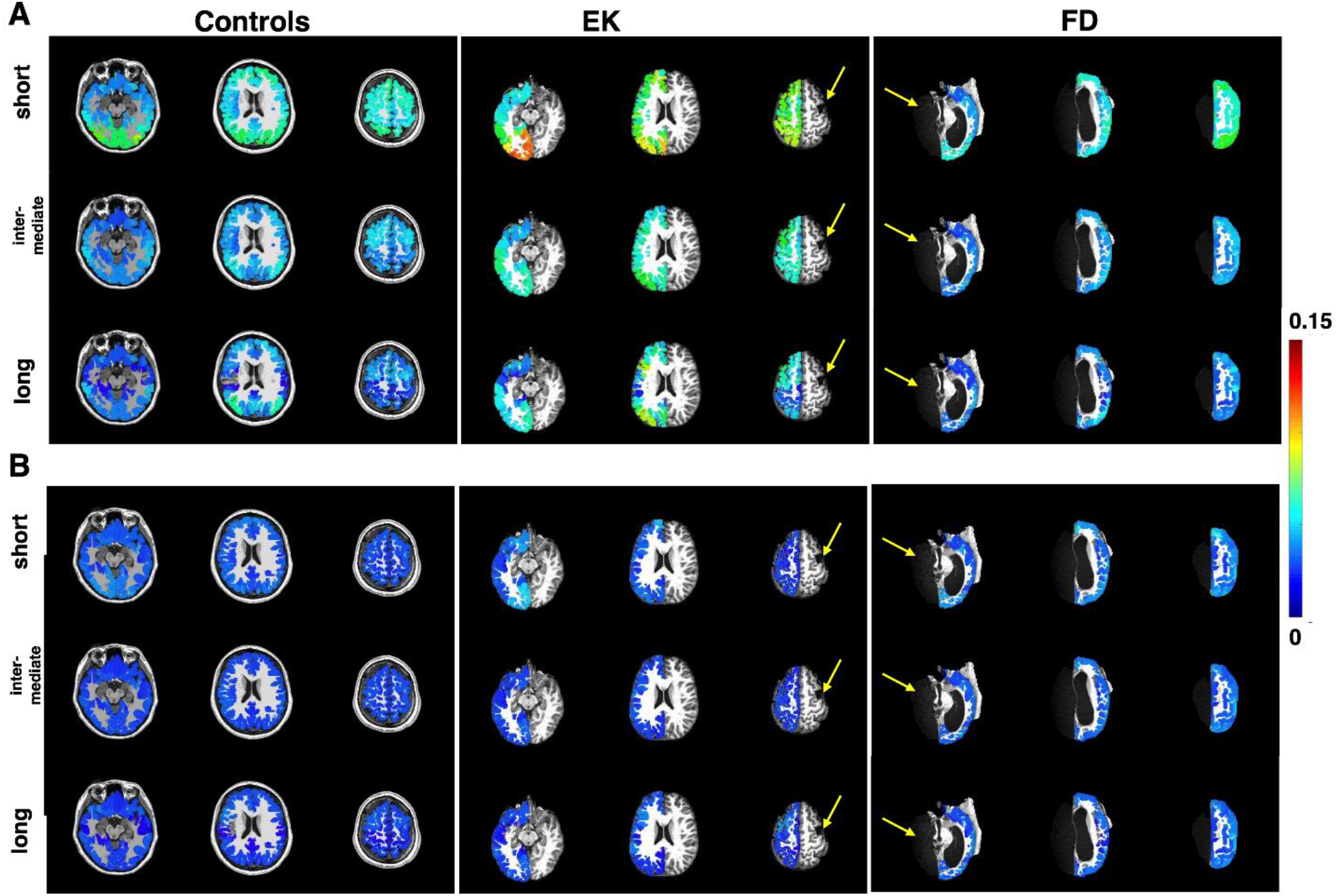
Mean significant connectivity fraction (SCF) in controls and two exemplar patients at different axial slices. Controls have low mean SCF with values not reaching 0.15 (highest at short-range) indicating more voxels cooperatively responsive at shorter than longer distances. In a patient’s contralesional hemisphere (e.g., LH in EK and RH in FD), SCF values are elevated, indicating a larger fraction of voxels with highly correlated time series across all distances (c.f. Table 2). Lesions in EK (due to a right focal resection) and FD (due to left hemispherectomy) are indicated by yellow arrows.

In contrast with controls, the positive-SCF in the contralesional hemisphere of patients such as EK were evidently different to controls (Fig. 5A, middle panel, patient’s SCF appear higher than controls’ SCF), while the negative-SCF in EK appeared comparable to controls (Fig. 5B, middle panel, all blue). Positive- and negative-SCF maps of all the other patients are included in the Supp. Mat. Fig. S8 and S9, respectively.

Next, we quantified the differences between the patients and controls at the group level. We used a paired Wilcoxon rank-sum test on the mean SCF in each of the 180 ROIs in each hemisphere, as defined in the HCP atlas. With this, we were able to determine whether the SCF were consistently different between a single patient and the mean in controls – that is, if the SCF were either consistently higher or consistently lower than corresponding values in controls (Table 2). We did this separately for the positive- and negative-SCF, and found that there were changes across all distances for both the positive- and the negative-SCF. Overall, this suggest an increase in the volume of cooperative voxels in the patients’ contralesional hemisphere.

**Table 2.** Median positive- and negative-SCF, in patients’ contralesional hemispheres and controls’ corresponding hemisphere for three different communities. Bold values indicate median SCF in patients that are larger than that in controls. Significance values from Wilcoxon rank-sum test comparing matched SCF between patients vs. controls with Bonferroni correction. *p<0.05, **p<0.01, ***p<0.001

| Hemisphere | Median, positive-SCF, Patients |  |  | Median, positive-SCF, Controls |  |  |
| --- | --- | --- | --- | --- | --- | --- |
|  | short | mid | long | short | mid | long |
| LH | **0.0613\*\*\*** | **0.0396\*\*\*** | **0.0233\*\*\*** | 0.0397 | 0.0207 | 0.0142 |
| RH | 0.0366 | 0.0195 | 0.0153 | 0.0355 | 0.0197 | 0.0132 |
| Hemisphere | Median, negative-SCF, Patients |  |  | Median, negative-SCF, Controls |  |  |
|  | short | mid | long | short | mid | long |
| LH | 0.0093 | 0.0077 | **0.0082\*** | 0.0103 | 0.0081 | 0.0072 |
| RH | **0.0122\*\*\*** | **0.0091\*\*\*** | **0.0088\*\*\*** | 0.0098 | 0.0077 | 0.0061 |

We also quantified the SCF on an individual level and found heterogeneity in the changes (Supp. Mat. Table S10). For some patients (EK and UD), the positive-SCF was significantly different to the controls for all three distances (median in patients > median in controls), while in others, the positive-SCF was different in only two (KQ, TC: median in patients > median in controls; NN: median in patient < median in controls) or in only one distance (DX: median in patient < median in controls). Still, three patients (SN, FD, JF) did not exhibit any significant differences in the positive-SCF compared to controls for any of the distances. For the negative-SCF, some patients (UD and FD) differed from the controls for all three distances (median in patients > median in controls), while in others, the negative-SCF was different in only two (EK, KQ: median in patients < median in controls) or in only one distance (JF: median in patient > median in controls). Interestingly, patient (NN) had an intermediate median negative-SCF that was smaller than controls’, and a long-range median negative-SCF that was larger than controls’. And finally, there were two patients (DX, TC) who did not exhibit altered negative-SCF. Interestingly, even though the negative-SCF of FD (Fig. 5B, right panel) were less visibly different to controls (c.f. Fig. 5A, EK versus controls), the values across all distances were nevertheless significantly different (note that we are only showing a few axial slices).

In sum, we have uncovered group-level increases with alterations among individual patients to the cortical volume that exhibited strongly correlated activation. Given that these patients exhibited largely normal visuoperceptual and/or cognitive abilities (except NN), our results might indicate that group differences in cortical volume reflect differential recruitment of cortex to mediate normal behavior. It is noteworthy that between this SCF analysis and the variable and stable network comparisons (Sections 2.2.2 and 2.2.3), almost all the patients evince enhanced negative correlations compared with the controls and this is somewhat more pronounced for those with a preserved right hemisphere.

### 2.4 Summary of analyses and results

Finally, and in summary, all of our analyses and the relevant results are summarized in Table 3. We compared FC at the functional and anatomical ROI level, and at the network level, in patients versus controls and found no group-level differences in either the between- or within-ROI/network FC. There were also no differences between controls and patients at the single-subject level (except for FC of functional ROIs in 2 patients only: NN and JF). Further, between-ROI/network was always weaker than within-ROI/network FC, indicating greater correlation among voxels from the same ROI/network.

**Table 3.** Summary of all analyses and results with relevant figures and tables for reference.

| Analysis | Results | Relevant Figures/Tables |
| --- | --- | --- |
| Connectivity of 7 functional ROIs | All voxels within the same functional cortical ROIs were positively correlated (no negative FC within-ROI) | Supp. Mat. Figure S2, Table S3 |
|  | Voxel-wise positive and negative FC (roughly 50%/50%) among subcortical ROIs |  |
|  | FC between < FC within the same ROIs |  |
|  | No group-level differences in patients and controls for between-ROI FC or within-ROI FC |  |
|  | Single subjects in control distribution (except NN and JF) |  |
| Connectivity of 180 anatomical ROIs | Voxel-wise positive and negative FC (roughly 50%/50%) among all pairs of ROIs | Figure 1, Supp. Mat. Table S4 |
|  | FC between < FC within ROIs |  |
|  | No group-level differences in patients and controls for between-ROI FC or within-ROI FC |  |
|  | No single-patient differences compared to control group |  |
| Connectivity of 22 anatomical networks | Voxel-wise positive and negative FC (roughly 50%/50%) among all pairs of networks | Figure 2, Supp. Mat. Table S5, Supp. Mat. Figure S6 |
|  | Negative voxel-wise FC was not an artefact of pre-processing |  |
|  | FC between < FC within the same networks |  |
|  | No group-level differences in patients and controls for between-ROI FC or within-ROI FC |  |
|  | No single-patient differences compared to control group |  |
| FC of isolated network pairs and individual subjects versus controls | Positive FC stable and normal in all patients vs controls | Figure 3 |
|  | KQ, UD, NN, FD, JF showed abnormal negative FC compared to controls |  |
| Stability of FC in network pairs | Most network pairs had high variability, while some had stable, equal FC across all controls | Figure 4 |
|  | Patients exhibited altered FC compared to controls in the supposedly stable network pairs |  |
|  | No single network pair is altered in all patients compared to controls, but two network pairs were altered in all three RH resection patients |  |
| Significant correlation fraction | Group level differences in both positive- and negative-SCF across all distances | Figure 5, Table 2, Supp. Mat. Figures S8, S9, Table S10 |
|  | Single-subject level differences, albeit a heterogenous mix, in all patients versus controls |  |

In all participants, be it controls or patients, we separated the voxel-wise FC into positive or negative correlations, which were present at roughly equal proportions among anatomical ROIs/networks. We confirmed that the negative FC was not simply a result of data pre-processing and, in fact, when comparing the individual corresponding network pairs in patients and controls, there were single-subject differences only in the negative FC of five patients compared to controls, while the positive FC was stable in all patients. Last, we introduced here a possible neuroimaging biomarker, the significant correlation fraction (SCF), which captures information about the cortical volume that shows high positive or negative correlation (i.e. proportional to highly correlated voxels). Whereas the SCF was typically low in controls, in a majority of the resection patients, the SCF was altered in the contralesional hemisphere. These changes were seen in the focal resection and hemispherectomy patients and hemisphere resected did not obviously explain these patterns.

## 3 Discussion

Surgical resection of epileptogenic tissue is not only a viable and effective treatment option for those with pharmacoresistant seizures, but it can also result in substantial positive gains in cognitive abilities, especially in children (Helmstaedter et al., 2020). To elucidate the neural substrate supporting these post-surgical changes, we compared the functional connectivity (FC; correlation of time series) from patients with focal cortical resection or hemispherectomy and typically developing controls. At the outset, we validated our data by showing that the correlations between functionally-defined ROIs in the visual system evince strong positive correlations equivalent to those we see in controls. This confirms that ROIs, which function collaboratively and subserve the same behavior, are positively co-activated.

We then undertook analyses spanning all of cortex and across a wide range of measures, of voxel-wise, network level and specific networks pairs for both positive and negative FC at the level of anatomically-defined ROIs. We also demonstrated in an assumption-free fashion, a measure of the extent of cooperation among voxels across short, intermediate and long-range distances. Converging evidence from these analyses uncover alterations to the contralesional hemisphere in pediatric patients following surgical intervention. Whereas positive FC was normal overall (and some showed increase in positive-SCF), the alterations were especially evident in the magnitude of the negative FC, for individual network pairs and stable network pairs as well as in the increase in negative-SCF. Across multiple measures, almost every patient shows some change in negative FC, although in some cases, this is true to a greater extent than in others. The altered negative correlations in patients is also notable as there are far fewer network pairs that show high variability (Section 2.2.2) in negative (LH: up to 75, RH: up to 49) than positive FC (LH: up to 158, RH: up to 168) in controls.

One surprising finding is that, in contrast with Kliemann et al. (2019) who reported increased connectivity in between-but not within-network correlations patients versus controls, we documented equal between- and within-ROI and between- and within-network correlations, and this was so for both positive and negative correlations. A perhaps obvious explanation relates to the ages of the patients: whereas Kliemann et al. (2019) examined adults (even though their surgery was in childhood), our patients are children or adolescents. These younger patients are likely still in the throes of developmental change and emergent neural sculpting and this involves both strengthening within- and between-region correlations. The establishment of connectivity and circuitry is complete in the adults and the differential within- and between-region correlations is already accomplished. Longitudinal investigation of our participants will permit adjudication of this difference.

### There is no default universal backup, rather, all available resources are utilized

While we only have nine patients, there was enough variation in the resection site to warrant speculation as to the presence of a “default universal backup” and our reasoning goes as follows: in the event of the resection in either hemisphere, would the homologous site in the opposite hemisphere compensate by assuming the cognitive load of the resected tissue? Alternatively, is there a specific cortical region that might be (quasi-) perpetually plastic and able to take over the function of any diseased and/or resected tissue? Neither of these hypotheses is supported by our data: some patients with LH resection that impacted their VOTC (NN, FD, JF) exhibited alterations to the FC in the homologous visual cortex while others, also with LH VOTC resection (SN, TC), and those with RH VOTC resection (KQ, UD) did not. Therefore, there are no obvious consistent alterations to regions homologous to the resected tissue. Furthermore, there was not a single network pair that was consistently altered in all the patients, thus precluding the idea of a universal backup.

Instead of a specific region or network being recruited for functional substitution (Anderson et al., 2011), our findings are more suggestive of an “all hands on deck” approach, exhibited in the widespread changes to varying degrees in the positive- and negative-SCF, which characterizes the interconnected voxel communities, and the marked alterations to the negative FC – remarkably, even in the presumably stable network pairs – in the contralesional hemisphere of patients. Together, these point towards a converging conclusion: all available resources are utilized, possibly via different mechanisms (change in magnitude, change in volume, or an aggregate), and this results in largely normal cognition in unilateral resection patients. Having stated this, however, the ubiquity of the negative correlations, which are very infrequently observed in controls, warrants further consideration.

### Alterations in negative FC in post-surgical cases

Conventionally (although there are counterexamples such as in Qian et al., 2018), FC has largely been quantified in terms of *mean* correlation values, *r*, among all time courses of interest. However, this does not take into account that *r* ranges from −1 to 1. These values cancel out to some degree, if both positives and negatives are present, even at unequal proportions, thus failing to capture potential dynamical changes to the strength of correlated activity following an insult to the brain. There has, however, been growing recognition of the relevance of the directionality of the correlation value. Some have argued that negative correlations are merely an artifact of a global regression procedure (Murphy et al., 2009), while others have claimed that correlated and anti-correlated FC jointly serve as the foundation for cognition (Fox et al., 2005, 2007) and that the negative FC reflects the operation of anti-correlated networks that occur naturally and spontaneously in cortex. These fluctuating intrinsic correlations, reflected in both positive and negative FC, have been linked with neuropsychological and neuropsychiatric disorders (Guo et al., 2016, 2017, Wang et al., 2018), and assumed to be the consequence of atypical functional circuits (Gabrieli & Ford, 2012).

Here, we observed that the patients with focal resection or a complete hemispherectomy have largely normal cognition post-surgically, and, concurrently, abnormal negative FC (with few exceptions). The key question is what role might be played by the overabundance of negative connections. We assume that complex cognitive functions are carried out by the coordinated – and thus correlated – activity of multiple brain regions. This coordination is aided to the extent that the regions are anatomically connected, which in turn is facilitated by co-localizing them within the same hemisphere. This gives rise to an organization in which related functions are more likely to be co-localized within a hemisphere (for example, language dominance and the visual word form area; Van der Haegen et al., 2012), and unrelated functions are more likely to be localized in different hemispheres. As a result, in the normal brain, within-hemisphere FC tends to be more positive, and between-hemisphere FC tends to be more negative (Toro et al., 2008).

In the case of reorganization following unilateral resection, functions – related and unrelated – may be primarily co-localized within a single hemisphere (even a focal resection can disrupt remote regions of the single hemisphere as in diaschisis). As a result, when compared to a normal hemisphere, the positive FCs are preserved, because related functions are still co-localized within the hemisphere. However, the negative FCs are increased, because now more unrelated functions are co-localized within the same hemisphere, as compared to the normal case. We suggest then that the negative connectivity in the preserved hemisphere plays the role of sculpting collaborative circuits by anti-correlating regions that are not engaged in the relevant behavior. There is one further desideratum of this plasticity – it has to be ready and in situ given that positive cognitive outcomes are evident even at one year post surgery (Helmstaedter et al., 2020). Any plasticity that might support this improvement, then, must be readily available, and the existing ubiquitous and intrinsic dynamic fluctuations amongst cortical regions (Fox et al., 2005, 2007) might play this key role.

If the negative correlations do play the role of anti-correlating unrelated regions or networks, this leads to the prediction that the less related the functions of ROIs or networks, the more evident their negative FC should be. In their early work, Fox et al (2005) showed anti-correlations between networks that are task-positive (activation) and task-negative (de-activation). Indeed, the idea of opposing or competing processes is well established, with negative interactions between, for example, focused attention versus general monitoring of one’s environment (Corbetta & Shulman, 2002) or between regions associated with cognition versus emotion (Simpson et al., 2001). In the current paper, the two network pairs that have most consistent deviation from control mean may be instances of unrelated functions, are the posterior cingulate cortex and somatosensory/motor cortex and the other is between posterior cingulate cortex and posterior opercular cortex. Close scrutiny of the map of cortical correlations in Fox et al. (Figure 3, 2005) shows exactly these networks in opposition (anti-correlated). Our findings suggest then that the very same anti-correlation ROIs/networks present in non-neurological cases are also present in the patients and potentially recruited on a much more significant scale post-surgically.

### Limitations of the study

Although we have presented converging evidence from a range of analyses, there are, nonetheless, obvious limitations. First, the parcellations used to create ROIs and networks in the patients’ native volumetric space were from the HCP atlas (Glasser et al., 2016). Glasser et al. (2016) encouraged registration of their atlas to individual subjects’ surface space, but this was not ideal in light of the (large) lesions in our patient cohort. While it is nevertheless possible to create surface files for the patients as was done in Kliemann et al. (2019), to facilitate a reproducible pipeline without the need for highly specialized manual interventions, we opted to analyze and use volumetric data, with the caveat that the atlas registration may not be absolutely perfect. To mitigate potential consequences, we were very conservative in our data pre-processing. We analyzed the time courses only of those voxels inside the grey matter mask of each participant. Additionally, we conducted other analyses that did not rely on the assumptions of the HCP atlas.

Second, volumetric distance is not the same as surface distance. In volumetric space, two nearby voxels may be on either side of a gyrus, in which case, surface distance (as in white matter connection) would actually be larger than the volumetric distance. However, we used a consistent processing pipeline for both patients and controls and any artefacts introduced by the difference in volumetric and surface distance would be systematic error, rather than random error, and should not affect our conclusions.

Last, as with many neuropsychological studies, we have a limited number of patients in this study. This makes it difficult to generalize our findings to the population of children who have undergone or will undergo resective surgery, and the scope for future investigations is broad. This notwithstanding, there are replicable profiles (e.g. distance effects, negative FC) across all patients largely independent of size and site of lesion, perhaps reflecting a robust and replicable algorithm across patients.

### Final remarks

Our results of altered negative FC between different networks and larger non-specific volume of cortex exhibiting highly correlated activity point to a theoretical increased cost for functional specificity in the limited cortical territory following a unilateral resection (either due to focal resection or hemispherectomy). If true, this would be intuitively consistent with the notion that functions (and behaviors) need to be represented in the typical brain; however, in a surgically resected brain, the cortical tissue is reduced, while the functions remained unchanged. Given that the patients have largely normal cognition and perception, it is possible that the cost might be metabolic in nature and does not manifest in neuropsychologically measurable gross deficits to behavior. Last, while the stable positive FC profiles in these patients might be sufficient, and perhaps the only relevant neural signature, in sustaining normal behaviors, the combination of changes we observed and especially the altered negative FC suggest, provocatively, that these too play a critical role in reorganization of function.

## 4 Materials and Methods

The procedures used here were reviewed and approved by the Institutional Review Boards of Carnegie Mellon University and the University of Pittsburgh. Parents provided informed consent and minor participants gave assent prior to the scanning sessions. Using the cortical parcellation based on the Human Connectome Project (HCP) atlas (Glasser et al., 2016), we analyzed the strength of connectivity of the contralesional hemisphere from visually evoked BOLD signals. The HCP atlas has 180 ROIs comprising 22 distinct networks in each hemisphere. We used these anatomically defined ROIs and networks, as well as functionally defined visual category ROIs in our analyses. Additionally, we present here a novel way of characterizing reorganization in terms of FC with minimal assumptions and based on linear distance alone.

All data and codes used in this study will be publicly available from KiltHub, Carnegie Mellon’s online data repository (doi: 10.1184/R1/12616316), upon publication.

### 4.1 Participants

Nine children or adolescents who had undergone unilateral surgical resection participated in this study (Table 1). Most of the patients underwent surgery at UPMC Children’s Hospital of Pittsburgh, Pittsburgh, USA. In addition, nine age-matched typically developing healthy control children and adolescents were also recruited (patients’ mean age 15.6 ± 1.5 years; controls’ mean age 13.7 ± 2.4 years; no significant difference between group ages, Wilcoxon rank sum test p > 0.08).

### 4.2 MRI parameters

Anatomical and functional images were acquired on a Siemens Verio 3T scanner with a 32-channel head coil at Carnegie Mellon University for most of the participants. However, data from one patient (JF) were acquired on a Siemens Prisma 3T scanner with a 64-channel head coil. The acquisition protocols were similar in the two sites.

For each participant, we obtained a T1-weighted anatomical image using the MPRAGE sequence (1mm isotropic resolution, TE=1.97 ms, TR=2300 ms, total scan time=5 minutes 21 seconds) and three sets of conventional visual category-localizer fMRI data (2mm isotropic resolution, TE=30 ms, TR=2000 ms). During the fMRI scans, participants watched images back-projected onto a screen mounted outside the bore and reflected by a mirror mounted on the head coil toward participants’ eyes. The visual stimulation paradigm, which is normally used to localize category-selective cortical regions, involved the presentation of 16-second blocks of images from different visual categories including faces, scenes, objects, words, and scrambled objects, interleaved with 8-second fixation blocks (scan time per run: 6 minutes 8 seconds). To ensure fixation, participants were asked to press a response button when the same image appeared consecutively in the stream (one-back task) and there was one pair of identical consecutive images per category. Control participants were placed in a simulator prior to the experimental data acquisition and were trained to lie still (receiving feedback when movement was detected). The patients have all had many scans as part of their medical management and had been trained previously to lie still.

### 4.3 Data pre-processing for FC analyses

All data were co-registered to the anatomical image and processed in each subject’s native volumetric space. fMRI BOLD data were pre-processed, using AFNI ‘Claudius’ v 19.2.26 (Cox, 1996). The AFNI package has been shown to have the most accurate motion estimation, as well as the least smoothing of data (Oakes et al., 2005) that worked well for children, who may especially be prone to movements inside the scanner. Pre-processing included, for each dataset, the following steps: all volume images were registered to the volume image with the least motion, and the aligned 4D series data were volume-registered to the participant’s skull-stripped anatomical image. Further, the time series of each voxel was despiked and corrected for slice-time acquisition offset. Motion in six directions (three for translation, three for rotation) and its mean and motion derivative, as well as the mean signal from the white matter voxels, were regressed out of each voxel’s time series. Time points with motion greater than 0.3 mm or with greater than 10% of voxels with outlier signal were censored to zero (minimal in controls, at most 12/184 points in a single run in patients). The time series after all these pre-processing steps were used in the FC analyses. In Supp. Mat. Fig. S6, we show the effects of various pre-processing pipelines on data quality.

### 4.4 Defining the functional category-specific ROIs

Image pre-processing included time-slice correction, volume registration, and co-registration. Functional images were volume-registered to the structural image with the least motion and outliers, and then co-registered to the skull-stripped anatomical image. Additionally, volumes with more than 0.3 mm motion and those with more than 10% voxel outliers were censored. The BOLD amplitude was also scaled to a uniform mean of 100 and uniform maximum of 200 arbitrary units. A general linear model (GLM) of the stimulation time course convolved with a canonical hemodynamic response function was used to determine category-selective voxels at p<0.001 and a cluster estimation algorithm was used to discard statistically insignificant smaller clusters of voxels at p<0.05. Last, we used anatomical masks from the HCP atlas to eliminate ROIs outside the VOTC.

Additionally, subcortical ROIs (LGN and pulvinar) were manually drawn on the anatomical image by EF and independently validated by SK.

### 4.5 Anatomical parcellation

In order to define the anatomical cortical regions, we used a script, provided to us by Dr Daniel Glen (Computer Engineer, NIH) that implemented AFNI commands to co-register the HCP atlas in MNI space to the native volumetric space for both groups of patients and controls. Briefly, each participant’s skull-stripped anatomical image was normalized to the HCP atlas in MNI space using affine and non-linear transformations, as well as dilations (to account for size differences between children’s brain and adult brains in HCP atlas). Next, a reverse transformation was applied to the HCP atlas to obtain a corresponding parcellation in native space. We used all 180 ROIs that together form 22 HCP-defined networks from each (or, in patients, the contralesional) hemisphere as the nodes in our connectivity analysis (see Supplementary Table S2).

We note that while the HCP atlas is ideally transformed to a single subject’s native space using surface registration, the challenges in producing the surface images of the patient’s structurally altered brain precluded us from doing so. In striving for minimal user intervention that might possibly introduce inconsistencies and in the hope that the pipeline we adopt here could be as highly reproducible as possible, we elected to analyze everything in native volumetric space to minimize the distortions – a consideration that is especially critical in cases where there are not only large areas of the cortex missing in the ipsilesional hemisphere, but in some cases, there are also accompanying morphological abnormalities even in the contralesional hemisphere as a result of, for example, a midline shift.

### 4.6 Connectivity of the contralesional hemisphere

To characterize the connectivity profiles, we used task-evoked BOLD signals instead of resting-state BOLD signal. This offered the advantage of avoiding the lower signal-to-noise ratios typically seen in resting-state data compared to evoked BOLD data and higher signal would permit us to see even relatively small changes in FC. We used the Pearson correlation coefficient as a proxy for FC between all voxel pairs from any ROI- or network pair. For example, given *N* voxels in ROI/network A and *M* voxels in ROI/network B, we computed *N*x*M* Pearson correlation coefficients between all unique pairwise voxel-to-voxel combinations and averaged over all these values to get the FC between the ROI/network pair of A and B. We also divided the *N*x*M* computations depending on whether the correlations were positive or negative, and then computed the positive or negative FC as the mean of only the positively correlated voxels or of only the negatively correlated voxels, respectively. With this, we were able to obtain the number of voxels out of *N*x*M* combinations that were positively or negatively correlated.

### 4.7 Characterization of distance from mean

We compared patients’ FC values in patients to controls under two criteria. First, we computed the distance-from-mean for each network pair by subtracting the FC value in controls from the patients and dividing this by the standard deviation in controls. Thus, we found network pairs that deviated from the control mean by two standard deviations or more (criterion 1). Next, we counted the number of network pairs that satisfied criterion 1 (either greater or smaller FC in patients than controls). To determine, on an individual level, which patient had significant alterations, we performed a leave-one-out analysis on the nine control data sets. To that end, we compared a single control to the remaining eight controls and counted the number of network pairs that were more than two standard deviations away from the artificial group mean of the eight controls. We repeated this for all controls and obtained a range of numbers that we used as our significance cutoff. For example, for positive FC of the left hemisphere in controls, there are up to 158 network pairs out of the possible 253 network pairs (including between- and within-network) that deviated between any single control and the rest of the control group. For a patient to have a significant distance-from-mean map, they would need to have more than 158 network pairs that exhibited a distance-from-mean of two or more standard deviations from the mean of all the controls (criterion 2). We note, however, that this test does not indicate *which* network pairs were altered, but only gives us a benchmark against which we can compare an individual patient’s degree of variability.

### 4.8 Heatmaps of network pairs susceptible to alterations

Next, we wanted to determine whether there were specific network pairs that were more susceptible to change, presumably as a result of the resective surgery, after accounting for individual differences as seen in controls. To that end, using the results of the leave-one-out test in controls from the previous section, we mapped all the network pairs in controls that exhibited differences between any individual control and the mean of the remaining controls. Any network pair for which no single control participant showed a deviation from the artificial mean was considered a stable network pair. We hypothesized that if a patient were to show a deviation in the mean FC for such stable network pairs, it would more likely be due to the surgery rather than due to some individual idiosyncrasy. We also generated a patient group heatmap by combining the individual patient heatmaps based on the side of resection.

### 4.9 Significant correlation fraction maps in participants

Here, we used an assumption-free approach to FC by defining three communities for any given voxel: short, intermediate, and long-range communities. To do so, we first computed the rounded up maximum Euclidean distance, D, between any two pairs of voxels from the same hemisphere. We then linearly divided D into four equal segments, such that you have d: {2, d1, d2, D} where 2 is the voxel size and thus the minimum separation between any pair of voxels. For any voxel, the short-range community of voxels are those within the distance of [2, d1) in mm, inclusive of 2. Correspondingly, its intermediate-range community are voxels within the distance of [d1, d2) in mm, inclusive of d1 and its long-range community are voxels within the distance of [d2, D) in mm, inclusive of d2. For any and all voxels, we computed the correlation of its time series with all the voxels in each of its communities. We then counted the number of correlations that were |r|>0.241 (p<0.001, df=182), separately for the positive and negative correlations, and divided this by total the number of voxels in each community. This number is what we called the significant correlation fraction or SCF, which can be from either the positive or negative correlations (positive- or negative-SCF, respectively).

## Acknowledgements

The authors would like to thank the participants and their families for taking part in this study. We also would like to thank Dr Daniel Glen for his help in implementing individual subject volumetric image registration to the HCP atlas, Ms Jennifer Monahan of Children’s Hospital for assistance in patient recruitment, and Mr Scott Kurdilla and Mr Mark Vignone for assistance in scanning the participants. We also thank Dr Avital Hahamy and Dr David Plaut for useful suggestions and conversations.

This research was supported by a grant from the National Eye Institute (NIH) RO1 EY0207018 to MP and CP and a grant from the National Institute of General Medical Sciences (NIH) T32 GM081760 to MCG. The content is solely the responsibility of the authors and does not necessarily represent the official views of the National Eye Institute, National Institute of General Medical Sciences, or the National Institutes of Health.

## Author Contributions

AMSM: conception and design; acquisition, analysis, and interpretation of data; created new code to use in work; wrote and revised manuscript; MCG: acquisition, analysis, and interpretation of data; created new code to use in work; revised manuscript; EF: analysis, and interpretation of data; created new code to use in work; revised manuscript; SK: analysis, and interpretation of data; revised manuscript; MAP: analysis, and interpretation of data; CP: contributed unpublished tools; recruited and managed patients; revised manuscript MB: conception and design; interpretation of data; wrote and revised manuscript.

The authors declare no competing interests.

