## Supplementary Materials for "All hands on deck: Large-scale (re)sculpting of cortical circuits in post-resection children"

**Table S1.** *IQ and visual perceptual measures of patients. Patients largely exhibited normal performance compared to controls in intermediate and high-level vision. Even in some patients who showed atypicality in some measure, all other measures were normal (e.g. abnormal contour integration and object matching RT in NN, but all other measures were normal). *p<0.05, **p<0.01, ***p<0.001 after correcting for multiple comparisons, per individual. For more details, see Liu, Freud et al., 2019 and Maallo et al., 2020.*

| Patient Code | IQ measures | Intermediate-level vision | | | High-level vision | | | |
|---|---|---|---|---|---|---|---|---|
| | | Contour integration | | Glass pattern | CFMT-C | | Object matching | |
| | | Threshold (±0 collinearity) | Threshold (±20 collinearity) | Threshold | Upright Faces (%) | Inverted faces (%) | Accuracy (%) | RT (ms) |
| EK | N/A | 54.01 | 75.51 | 54.17 | 88.33 | 81.67 | 100.00 | 1192.12 |
| KQ | Presurgical: 76 | 68.45 | 74.21 | 35 | 90.00 | 78.33 | 99.00 | 1018.14 |
| UD | Presurgical: 116<br>Postsurgical: 118 | 51.87 | 76.83 | 25.83 | 83.33 | 68.33 | 91.00 | 1366.96 |
| DX | Presurgical: 86 | 69.24 | 78.17 | 36.67 | 100.00 | 93.33 | 96.00 | 1202.30 |
| NN | Presurgical: 67 | **75.95*** | 78.63 | 25.83 | 80.56 | 45.83 | 93.00 | **1726.1*** |
| SN | N/A | 64.61 | 78.17 | 33.33 | 95.00 | 73.33 | 95.00 | 897.31 |
| TC | N/A | 66.12 | 77.27 | 45.83 | 83.33 | 46.67 | 89.00 | 1047.66 |
| FD | N/A | 53.70 | 75.51 | 29.17 | 91.67 | 86.67 | 95.00 | **2274.13**** |
| JF | Postsurgical: 68 (DAS-II) | **78.17*** | 78.17 | 52.50 | 61.67 | 40.00 | 90.00 | 1491.44 |
| Control mean | | 56.45±5.05 | 74.04±3.53 | 39.4±7.74 | 88.22±11.55 | 70.36±11.79 | 94.86±3.08 | 869.03±225.65 |

**Figure S2.** *FC in visually-responsive regions in controls' left and right hemispheres and patients' contralesional hemispheres. (A, E, I, M) FC between seven visually-functionally-selective and anatomically-demarcated (LGN, Pulvinar) regions. (B, F, J, N) Negative FC fraction showing that between functionally specific regions, there are fewer negatively correlated voxels than between anatomically defined regions (e.g. compare blue EVC (early visual cortex) to EVC and green pulvinar to pulvinar). Nevertheless, there are still voxels that exhibit negative FC between functionally-specific regions, albeit at a weaker magnitude. Mean of voxels with only (C, G, K, O) positive correlations or (D, H, L, P) negative correlations. There are no within-region negative FC because within a region, all voxels were positively correlated. Note the stronger FC between functionally specific regions than anatomically-defined regions (c.f. Figs. 1 and 2). For some controls, we could not locate: word-selective region in the RH (6/9), LGN in the RH (1/9). Additionally, for some patients, we could not locate: word-selective region in the contralesional LH (2/3), object selective region in the contralesional LH (1/3), object-selective region in the contralesional RH (1/6). Means are shown from available data only.*

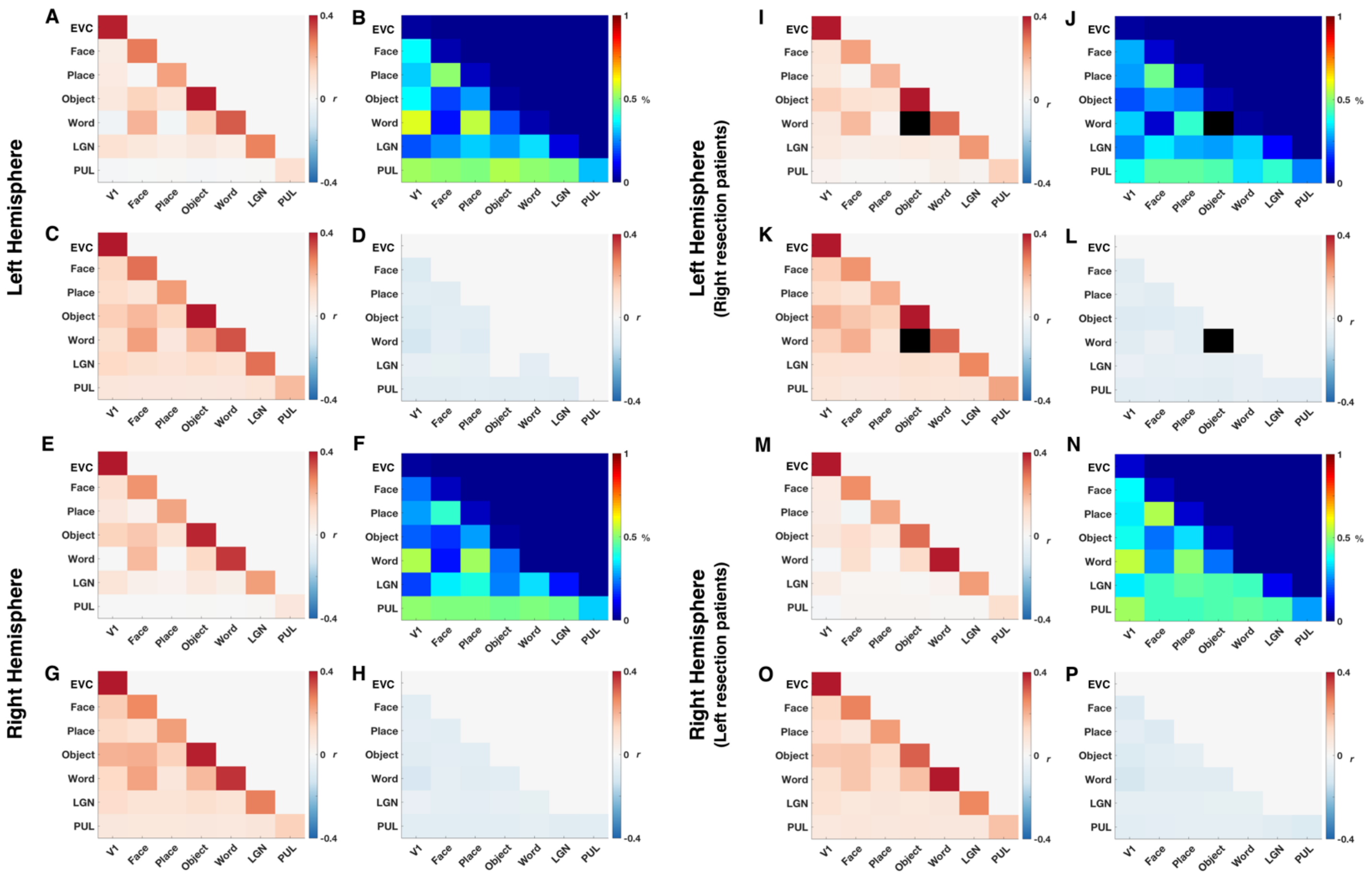

**Table S3.** *Comparison of within- and between-category ROI FC. Values between individual patient's contralesional hemisphere and controls' corresponding hemisphere were compared using modified t-test (Crawford & Howell, 1998). IndexFC is ratio of between-ROI to within-ROI FC (Kliemann et al., 2019). * p<0.05, ** p<0.01, ***p<0.001, after correcting for multiple comparisons (within- versus between- versus index, for mean FC, positive FC, negative FC) per individual.*

| | Participant | Within-network | Between-network | IndexFC |
|---|---|---|---|---|
| **Category-selective ROIs - Mean FC** | **Controls - LH mean** | 0.2843 | 0.0487 | 0.1686 |
| | **Controls - LH sdev** | 0.0382 | 0.0190 | 0.0556 |
| | **EK** | 0.2807 | 0.0599 | 0.2134 |
| | **KQ** | 0.2724 | 0.0598 | 0.2195 |
| | **UD** | 0.2472 | 0.0360 | 0.1456 |
| | **Controls - RH mean** | 0.2721 | 0.0547 | 0.2011 |
| | **Controls - RH sdev** | 0.0182 | 0.0131 | 0.0477 |
| | **DX** | 0.3186 | 0.0289 | 0.0907 |
| | **NN** | **0.1932*** | 0.0310 | 0.1605 |
| | **SN** | 0.2518 | 0.0467 | 0.1855 |
| | **TC** | 0.3099 | 0.0570 | 0.1839 |
| | **FD** | 0.2866 | 0.0581 | 0.2027 |
| | **JF** | 0.2506 | **-0.0102*** | **-0.0407**** |
| **Category-selective ROIs - Positive FC** | **Controls - LH mean** | 0.3063 | 0.1162 | 0.3810 |
| | **Controls - LH sdev** | 0.0343 | 0.0118 | 0.0332 |
| | **EK** | 0.3034 | 0.1075 | 0.3543 |
| | **KQ** | 0.2943 | 0.1111 | 0.3775 |
| | **UD** | 0.2801 | 0.1126 | 0.4020 |
| | **Controls - RH mean** | 0.2957 | 0.1145 | 0.3877 |
| | **Controls - RH sdev** | 0.0183 | 0.0105 | 0.0324 |
| | **DX** | 0.3446 | 0.1035 | 0.3003 |
| | **NN** | 0.2367 | 0.0992 | 0.4191 |
| | **SN** | 0.2710 | 0.1075 | 0.3967 |
| | **TC** | 0.3196 | 0.1142 | 0.3573 |
| | **FD** | 0.3098 | 0.1345 | 0.4342 |
| | **JF** | 0.3064 | 0.0808 | 0.2637 |
| **Category-selective ROIs - Negative FC** | **Controls - LH mean** | not applicable | -0.0752 | not applicable |
| | **Controls - LH sdev** | | 0.0088 | |
| | **EK** | | -0.0602 | |
| | **KQ** | | -0.0614 | |
| | **UD** | | -0.0857 | |
| | **Controls - RH mean** | | -0.0695 | |
| | **Controls - RH sdev** | | 0.0031 | |
| | **DX** | | -0.0764 | |
| | **NN** | | -0.0746 | |
| | **SN** | | -0.0639 | |
| | **TC** | | -0.0682 | |
| | **FD** | | **-0.1267***** | |
| | **JF** | | **-0.0887**** | |

**Table S4.** *Comparison of within- and between- anatomical ROI FC. Values between individual patient's contralesional hemisphere and controls' corresponding hemisphere were compared using modified t-test (Crawford & Howell, 1998). IndexFC is ratio of between-ROI to within-ROI connectivity (Kliemann et al., 2019). There were no significant differences between any individual patient and controls, after correcting for multiple comparisons (within- versus between- versus index, for mean FC, positive FC, negative FC) per individual.*

| | Participant | Within-network | Between-network | IndexFC |
|---|---|---|---|---|
| 180 ROIs - Mean FC | Controls - LH mean | 0.1326 | 0.0236 | 0.1742 |
| | Controls - LH sdev | 0.0298 | 0.0093 | 0.0386 |
| | EK | 0.1730 | 0.0530 | 0.3064 |
| | KQ | 0.1432 | 0.0412 | 0.2877 |
| | UD | 0.1386 | 0.0345 | 0.2489 |
| | Controls - RH mean | 0.1247 | 0.0215 | 0.1688 |
| | Controls - RH sdev | 0.0292 | 0.0095 | 0.0473 |
| | DX | 0.1332 | 0.0190 | 0.1426 |
| | NN | 0.0841 | 0.0247 | 0.2937 |
| | SN | 0.1309 | 0.0176 | 0.1345 |
| | TC | 0.1345 | 0.0209 | 0.1554 |
| | FD | 0.1120 | 0.0085 | 0.0759 |
| | JF | 0.1316 | 0.0133 | 0.1011 |
| 180 ROIs - Positive FC | Controls - LH mean | 0.1919 | 0.0971 | 0.5095 |
| | Controls - LH sdev | 0.0253 | 0.0083 | 0.0391 |
| | EK | 0.2236 | 0.1159 | 0.5183 |
| | KQ | 0.1937 | 0.1034 | 0.5338 |
| | UD | 0.2017 | 0.1112 | 0.5513 |
| | Controls - RH mean | 0.1861 | 0.0954 | 0.5162 |
| | Controls - RH sdev | 0.0244 | 0.0077 | 0.0382 |
| | DX | 0.1923 | 0.0940 | 0.4888 |
| | NN | 0.1606 | 0.1004 | 0.6252 |
| | SN | 0.1873 | 0.0925 | 0.4939 |
| | TC | 0.1955 | 0.0949 | 0.4854 |
| | FD | 0.1899 | 0.0925 | 0.4871 |
| | JF | 0.1956 | 0.0927 | 0.4739 |
| 180 ROIs - Negative FC | Controls - LH mean | -0.0818 | -0.0781 | 0.9573 |
| | Controls - LH sdev | 0.0065 | 0.0036 | 0.0358 |
| | EK | -0.0887 | -0.0750 | 0.8455 |
| | KQ | -0.0742 | -0.0704 | 0.9488 |
| | UD | -0.0945 | -0.0853 | 0.9026 |
| | Controls - RH mean | -0.0809 | -0.0778 | 0.9638 |
| | Controls - RH sdev | 0.0048 | 0.0024 | 0.0393 |
| | DX | -0.0831 | -0.0785 | 0.9446 |
| | NN | -0.0905 | -0.0814 | 0.8994 |
| | SN | -0.0797 | -0.0783 | 0.9824 |
| | TC | -0.0844 | -0.0772 | 0.9147 |
| | FD | -0.0915 | -0.0846 | 0.9246 |
| | JF | -0.0770 | -0.0807 | 1.0481 |

**Table S5**. *Networks used as nodes to compute the functional connectivity. These 22 networks were defined in the Human Connectome Project's multi-modal anatomical parcellation (Glasser et al., 2016).*

| **Label** | **Name** |
|---|---|
| *1* | Primary visual cortex |
| *2* | Early visual cortex |
| *3* | Dorsal visual stream |
| *4* | Ventral stream visual cortex |
| *5* | MT+ Complex and Neighboring Visual Areas |
| *6* | Somatosensory and Motor Cortex |
| *7* | Paracentral lobular and mid cingulate cortex |
| *8* | Premotor Cortex |
| *9* | Posterior opercular cortex |
| *10* | Early Auditory cortex |
| *11* | Auditory association cortex |
| *12* | Insular and frontal opercular cortex |
| *13* | Medial temporal cortex |
| *14* | Lateral temporal cortex |
| *15* | Temporo-parieto-occipital junction |
| *16* | Superior parietal cortex |
| *17* | Inferior parietal cortex |
| *18* | Posterior Cingulate cortex |
| *19* | Anterior cingulate and medial prefrontal cortex |
| *20* | Orbital and polar frontal cortex |
| *21* | Inferior frontal cortex |
| *22* | Dorsolateral prefrontal cortex |

**Figure S6.** *Effects of pre-processing pipeline. We compared the effects of various preprocessing pipeline on the positive/negative voxel population split from the data of a random control subject's LH. We regressed mean signal from (1-D)ventricles only or (E-H) white matter and ventricles. (I-L) We also looked at minimally processed volume-registered data, from which no nuisance signals were regressed. (M-P) Last, we also looked at the effects of regressing global mean signal in addition to signals from white matter and ventricles. Even minimally processed data showed negative voxel-wise correlations, albeit at smaller fractions than the other pre-processing pipelines, highlighting the need to address positive and negative FC separately.*

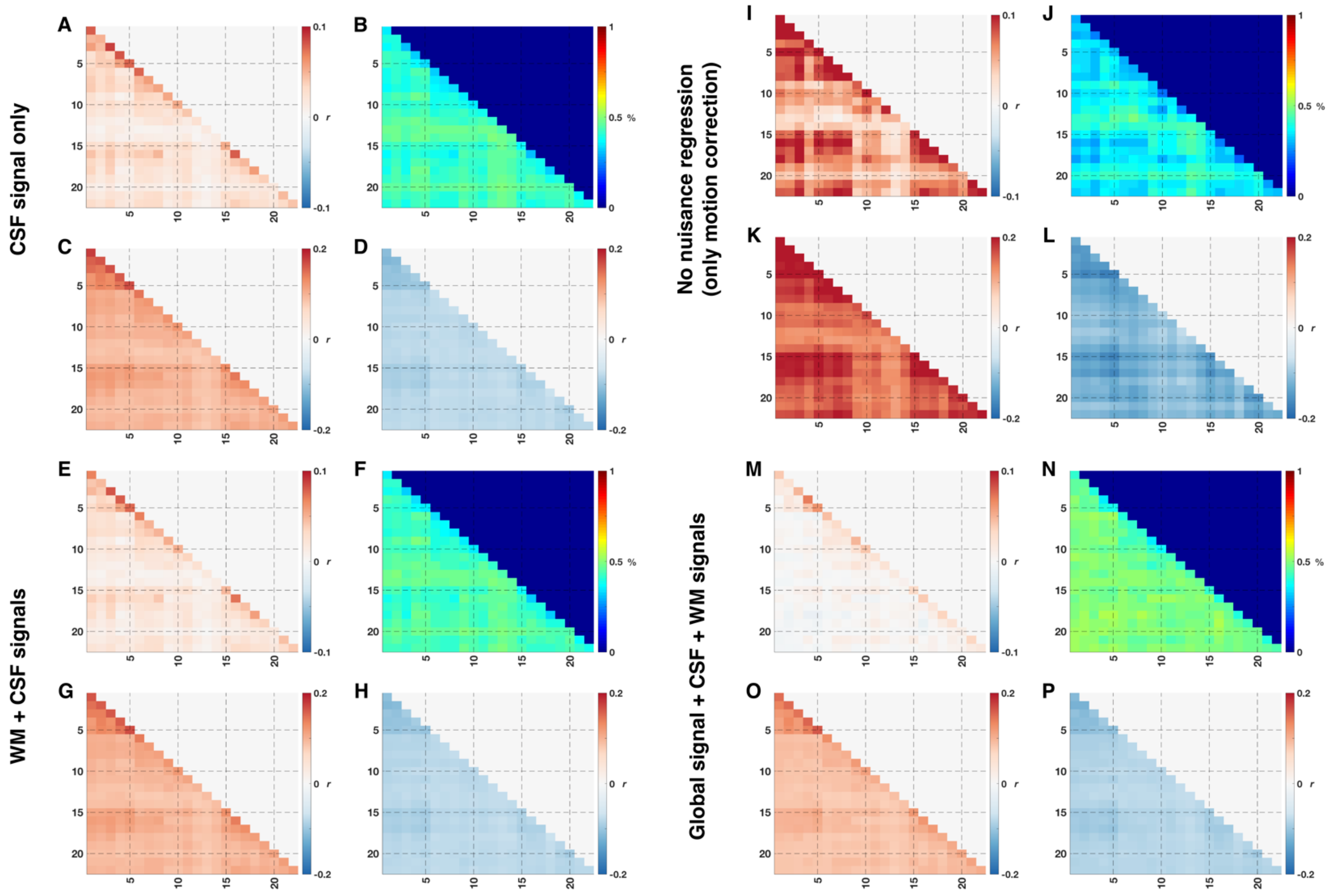

**Table S7**. *Comparison of within- and between-network FC. Values between individual patient's contralesional hemisphere and controls' corresponding hemisphere were compared using modified t-test (Crawford & Howell, 1998). IndexFC is ratio of between-network to within-network connectivity (Kliemann et al., 2019). There were no significant differences between any individual patient and controls, after correcting for multiple comparisons (within- versus between- versus index, for mean FC, positive FC, negative FC) per individual.*

| | Participant | Within-network | Between-network | IndexFC |
|---|---|---|---|---|
| Network - Mean FC | Controls - LH mean | 0.0640 | 0.0217 | 0.3310 |
| | Controls - LH sdev | 0.0146 | 0.0086 | 0.0703 |
| | EK | 0.0999 | 0.0516 | 0.5165 |
| | KQ | 0.0807 | 0.0379 | 0.4696 |
| | UD | 0.0769 | 0.0357 | 0.4642 |
| | Controls - RH mean | 0.0607 | 0.0213 | 0.3304 |
| | Controls - RH sdev | 0.0140 | 0.0093 | 0.0814 |
| | DX | 0.0604 | 0.0170 | 0.2815 |
| | NN | 0.0480 | 0.0243 | 0.5063 |
| | SN | 0.0621 | 0.0162 | 0.2609 |
| | TC | 0.0603 | 0.0195 | 0.3234 |
| | FD | 0.0380 | 0.0076 | 0.2000 |
| | JF | 0.0584 | 0.0119 | 0.2038 |
| Network - Positive FC | Controls - LH mean | 0.1391 | 0.0965 | 0.6953 |
| | Controls - LH sdev | 0.0131 | 0.0075 | 0.0323 |
| | EK | 0.1680 | 0.1170 | 0.6964 |
| | KQ | 0.1424 | 0.1008 | 0.7079 |
| | UD | 0.1528 | 0.1135 | 0.7428 |
| | Controls - RH mean | 0.1357 | 0.0950 | 0.7015 |
| | Controls - RH sdev | 0.0124 | 0.0068 | 0.0303 |
| | DX | 0.1392 | 0.0934 | 0.6710 |
| | NN | 0.1300 | 0.1005 | 0.7731 |
| | SN | 0.1360 | 0.0926 | 0.6809 |
| | TC | 0.1415 | 0.0960 | 0.6784 |
| | FD | 0.1276 | 0.0907 | 0.7108 |
| | JF | 0.1375 | 0.0919 | 0.6684 |
| Network - Negative FC | Controls - LH mean | -0.0839 | -0.0788 | 0.9395 |
| | Controls - LH sdev | 0.0042 | 0.0034 | 0.0146 |
| | EK | -0.0856 | -0.0764 | 0.8925 |
| | KQ | -0.0741 | -0.0707 | 0.9541 |
| | UD | -0.0907 | -0.0861 | 0.9493 |
| | Controls - RH mean | -0.0828 | -0.0780 | 0.9426 |
| | Controls - RH sdev | 0.0029 | 0.0019 | 0.0167 |
| | DX | -0.0852 | -0.0789 | 0.9261 |
| | NN | -0.0910 | -0.0817 | 0.8978 |
| | SN | -0.0816 | -0.0789 | 0.9669 |
| | TC | -0.0868 | -0.0793 | 0.9136 |
| | FD | -0.0901 | -0.0835 | 0.9267 |
| | JF | -0.0844 | -0.0810 | 0.9597 |

**Figure S8.** *Visualization of the positive-SCF in patients' contralesional hemispheres. Complementary data to Fig. 5. Lesions are indicated by yellow arrows.*

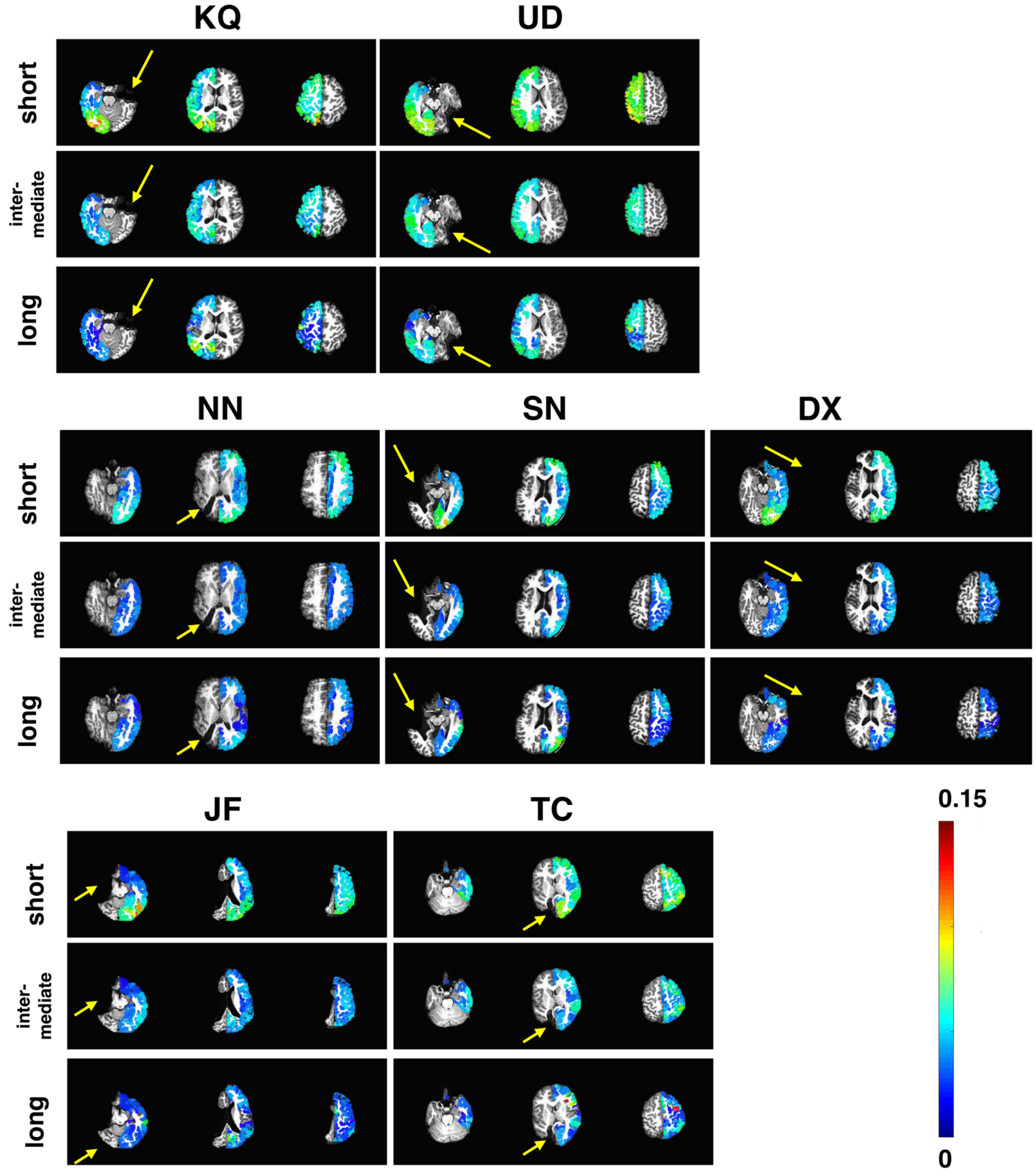

**Figure S9.** *Visualization of the negative-SCF in patients' contralesional hemispheres. Complementary data to Fig. 5. Lesions are indicated by yellow arrows.*

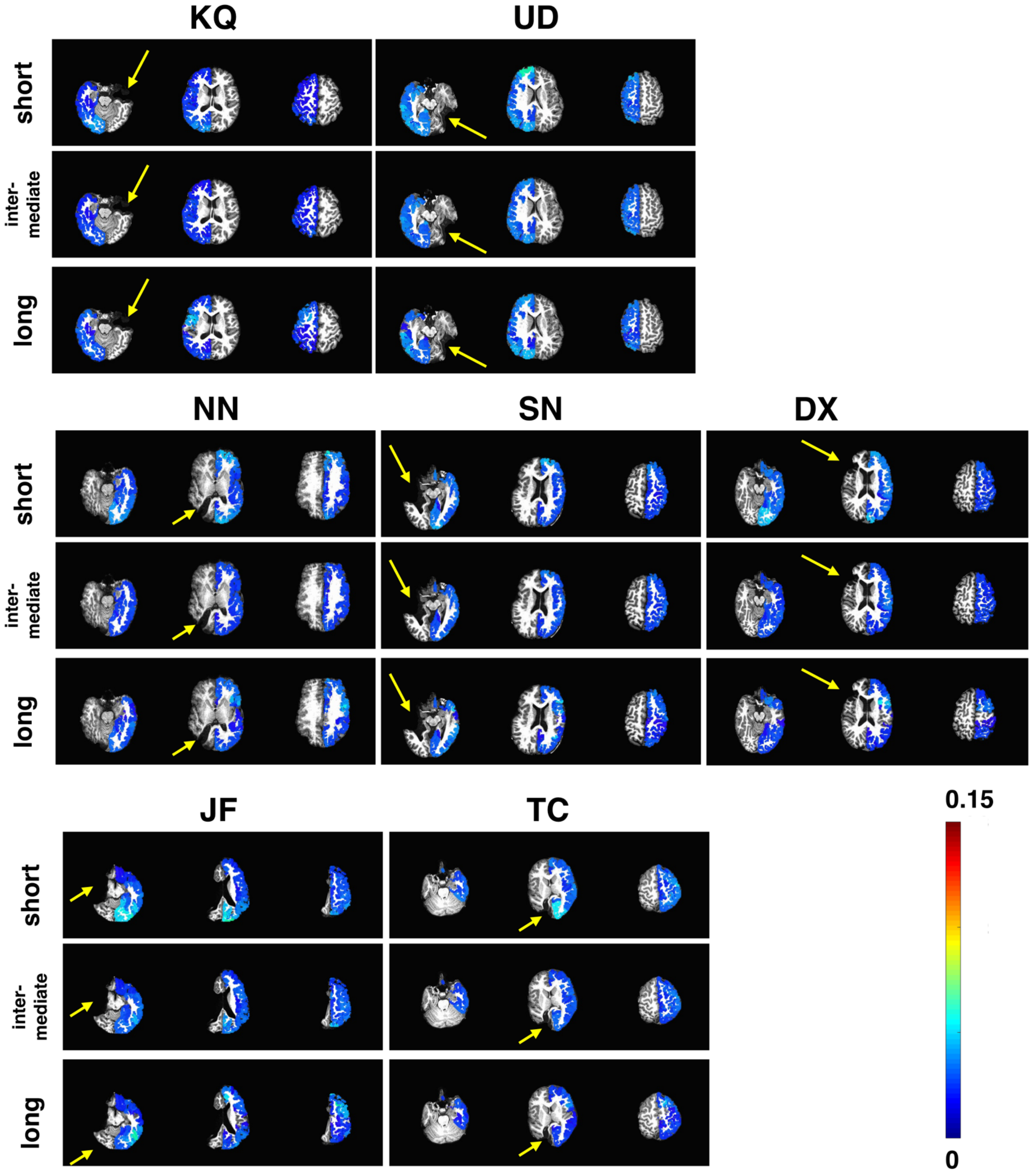

**Table S10**. *Median positive- and negative-SCF, in patients' contralesional hemispheres and controls' corresponding hemisphere for the three different communities. Values in bold (italics) indicate median SCF in patients that are larger (smaller) than median SCF in controls. Significance values from Wilcoxon rank-sum test comparing matched SCF between patients and controls with Bonferroni correction (multiplied by six, per individual). *p<0.05, **p<0.01, ***p<0.001*

<table>
<tr><th rowspan="2">Patient Code</th><th colspan="3">Median, positive SCF</th><th colspan="3">Median, positive SCF, Controls</th><th></th></tr>
<tr><th>short</th><th>mid</th><th>long</th><th>short</th><th>mid</th><th>long</th><th></th></tr>
<tr><td>EK</td><td>**0.0671*****</td><td>**0.0443*****</td><td>**0.0249*****</td><td rowspan="3">0.0397</td><td rowspan="3">0.0207</td><td rowspan="3">0.0142</td><td rowspan="3">LH</td></tr>
<tr><td>KQ</td><td>**0.0476*****</td><td>**0.0306*****</td><td>0.0159</td></tr>
<tr><td>UD</td><td>**0.0628*****</td><td>**0.0403*****</td><td>**0.0279*****</td></tr>
<tr><td>DX</td><td>0.0332</td><td>*0.0163***</td><td>0.0108</td><td rowspan="6">0.0355</td><td rowspan="6">0.0197</td><td rowspan="6">0.0132</td><td rowspan="6">RH</td></tr>
<tr><td>NN</td><td>*0.0296****</td><td>*0.0149****</td><td>0.0134</td></tr>
<tr><td>SN</td><td>0.0295</td><td>0.0177</td><td>0.0139</td></tr>
<tr><td>TC</td><td>**0.0444***</td><td>**0.0229***</td><td>0.0101</td></tr>
<tr><td>FD</td><td>0.0369</td><td>0.0206</td><td>0.0152</td></tr>
<tr><td>JF</td><td>0.0333</td><td>0.0191</td><td>0.0101</td></tr>
<tr><th rowspan="2">Patient Code</th><th colspan="3">Median, negative SCF</th><th colspan="3">Median, negative SCF, Controls</th><th></th></tr>
<tr><th>short</th><th>mid</th><th>long</th><th>short</th><th>mid</th><th>long</th><th></th></tr>
<tr><td>EK</td><td>*0.0083****</td><td>*0.0058****</td><td>0.0053</td><td rowspan="3">0.0103</td><td rowspan="3">0.0081</td><td rowspan="3">0.0072</td><td rowspan="3">LH</td></tr>
<tr><td>KQ</td><td>*0.0044****</td><td>*0.0043****</td><td>0.0053</td></tr>
<tr><td>UD</td><td>**0.0137*****</td><td>**0.0128*****</td><td>**0.0104*****</td></tr>
<tr><td>DX</td><td>0.0105</td><td>0.0074</td><td>0.0052</td><td rowspan="6">0.0098</td><td rowspan="6">0.0077</td><td rowspan="6">0.0061</td><td rowspan="6">RH</td></tr>
<tr><td>NN</td><td>0.009</td><td>*0.0062****</td><td>**0.0085*****</td></tr>
<tr><td>SN</td><td>0.0097</td><td>**0.0086****</td><td>**0.0080****</td></tr>
<tr><td>TC</td><td>0.0107</td><td>0.0079</td><td>0.0051</td></tr>
<tr><td>FD</td><td>**0.0144*****</td><td>**0.0121*****</td><td>**0.0115*****</td></tr>
<tr><td>JF</td><td>0.0097</td><td>**0.0096*****</td><td>0.0062</td></tr>
</table>
